# Myosin II independent contraction of actin filaments in membrane nanotubes

**DOI:** 10.1101/2025.09.04.674176

**Authors:** Md Arsalan Ashraf, Tapas Singha, Susav Pradhan, Serene Rose David, Pierre Sens, Pramod Pullarkat

## Abstract

The ability to generate active stresses within filamentous actin matrices is a fundamental and evolutionarily conserved process driving locomotion and morphogenetic changes in cells. The generation of pushing forces by actin polymerization is reasonably well understood, and is known to drive lamellipodia based motility and filopodial extension. Actin filaments decorated with myosin motors can also generate contractile stresses as in the cell cortex or in cytokinetic rings. In this article we use membrane nanotubes pulled out of axonal shaft to investigate actin dynamics and force generation. We report cyclic growth and retraction dynamics of actin within the tube and correlated contraction events giving rise to sustained load and fail cycles. The contraction mechanism operate independent of myosin II motor proteins. Furthermore, we analyzed the dynamics of actin within the tube, including under various biochemical or genetic perturbations. By combining these results with physical modeling, we argue that stresses generated in the actin filaments by the binding of actin depolymerizing factor (ADF/cofilin) proteins can explain the cyclic load-fail behavior.

## INTRODUCTION

Membrane nanotubes are generated by various cell types to perform important biological processes such as cell locomotion, environment sensing, signaling, intercellular transport, etc. Examples include filopodia generated by motile cells and neuronal growth cones [1], microvilli [2], steriocilia [3], pseudopodia of dendritic cells [4], and tunelling nanotubes that bridge multiple cells [5, 6]. The formation and stability of these nanotubes, and in some cases their growth and retraction dynamics, relies on the dynamics of actin filaments and their regulation by actin associated proteins. It is reasonably well established that such nanotubes can be formed by forces generated due to sustained polymerization of actin filaments [7, 8]. However, how actin dynamics is regulated within such nanotubes, for example, to drive growth and retraction of filopodia, and how some of structures like filopodia generate pulling forces, are still poorly understood, although several mechanisms have been proposed [9–12].

Here we show that membrane tethers drawn out of axons using optically trapped micron-size beads form a very convenient system to investigate actin dynamics within nanotubes. We demonstrate that actin filaments invade these extracted tubes and exhibit sustained growth and retraction cycles within the tube. More remarkably, such tubes exhibit actin-dependent contractile force generation via a mechanism which is independent of myosin II molecular motors. The observed cyclic load and fail behavior is quantified and compared across different biochemical and genetic perturbations to elucidate the mechanism responsible for the observed actin dynamics and force generation within nanotubes. Based on these, and a theoretical model, we propose a novel ADF/cofilin based mechanism that can explain the cyclic dynamics and the load-fail behavior in nanotubes.

The statistics of contractile stress generation is quantitatively reproduced by a model where multiple actin filament stochastically bind and unbind from the protrusion’s tip while undergoing continuous shrinkage due to cofilin binding.

### EXTRACTED NANOTUBES EXHIBIT LOAD AND FAIL CYCLES

Membrane nanotubes, also known as tethers, were extracted from axonal shafts of chick dorsal root ganglia neurons using an optically trapped polystyrene bead (Fig. 1a-c). After extraction of the nanotube, the distance between the axonal shaft (base) and the trap was maintained at a length of about 10 µm and the force on the bead was recorded. In many cases (≳ 50%), after about a minute, a repeating load and fail behavior (active force peaks) like the one shown in Fig. 1d could be observed (more examples in Suppl. Mat. Fig. S1). In the rest of the cases, where the tethers remained attached, no active force peak could be observed up to ≃10 min. When load-fail cycles were observed, the tether force increases slowly, pulling the bead towards the axon (see Fig. 1a), and eventually drops rapidly to a base value. The time evolution of the force build-up varied from cycle to cycle, but whenever the force collapsed back to the base value, the sudden drop in force exhibited a double-exponential decay (see Fig. S2). Histograms for the distributions of “rupture time” (*t*_rupt_), defined as the duration from start of a force event to final collapse to the base value (green horizontal line in Fig. 1d), and the peak force (*f*_peak_) within each cycle as measured from the baseline (blue vertical line in Fig. 1d), are shown in Figs. 1 e,f. To explore the mechanism for the active force generation, we applied various biochemical perturbations to the myosin motor proteins prior to nanotube extraction, and the results are shown in Fig. 2. No significant changes in the load-fail behavior could be observed after treatments using either the myosin II inhibitor Blebbistatin (20 µM), or with the Rho kinase inhibitor Y-27632, or with the myosin V blocker MyoVin-1 (a combination of MLCK inhibitor ML7 and Rho kinase inhibitor Y-27632 was also tried; see Suppl. Mat. Fig. S3). However, depletion of ATP (see Materials and Methods), stabilization of actin using Jasplakinolide, or prevention of actin polymerization using Latrunculin A completely abolished the load-fail cycles without exemption. These results demonstrate that the repeated occurrence of active force peaks require dynamic actin filaments but molecular motors like myosin II and myosin V are dispensable. As filament turnover seems to be involved, we also tried inhibiting actin nucleators: formins using FH2 domain inhibitor SMIFH2 and Arp2/3 using CK666. These inhibitors failed to abolish the active force peaks (see Fig. 2 and Suppl. Mat. Fig. S4). It has to be noted that the anti-formin drug SMIFH2 has been shown to also inhibit various myosins as well [13]. This include myosin IIA, Drosophila myosins V & VIIa, and bovine myosin X.

**Figure 1.**
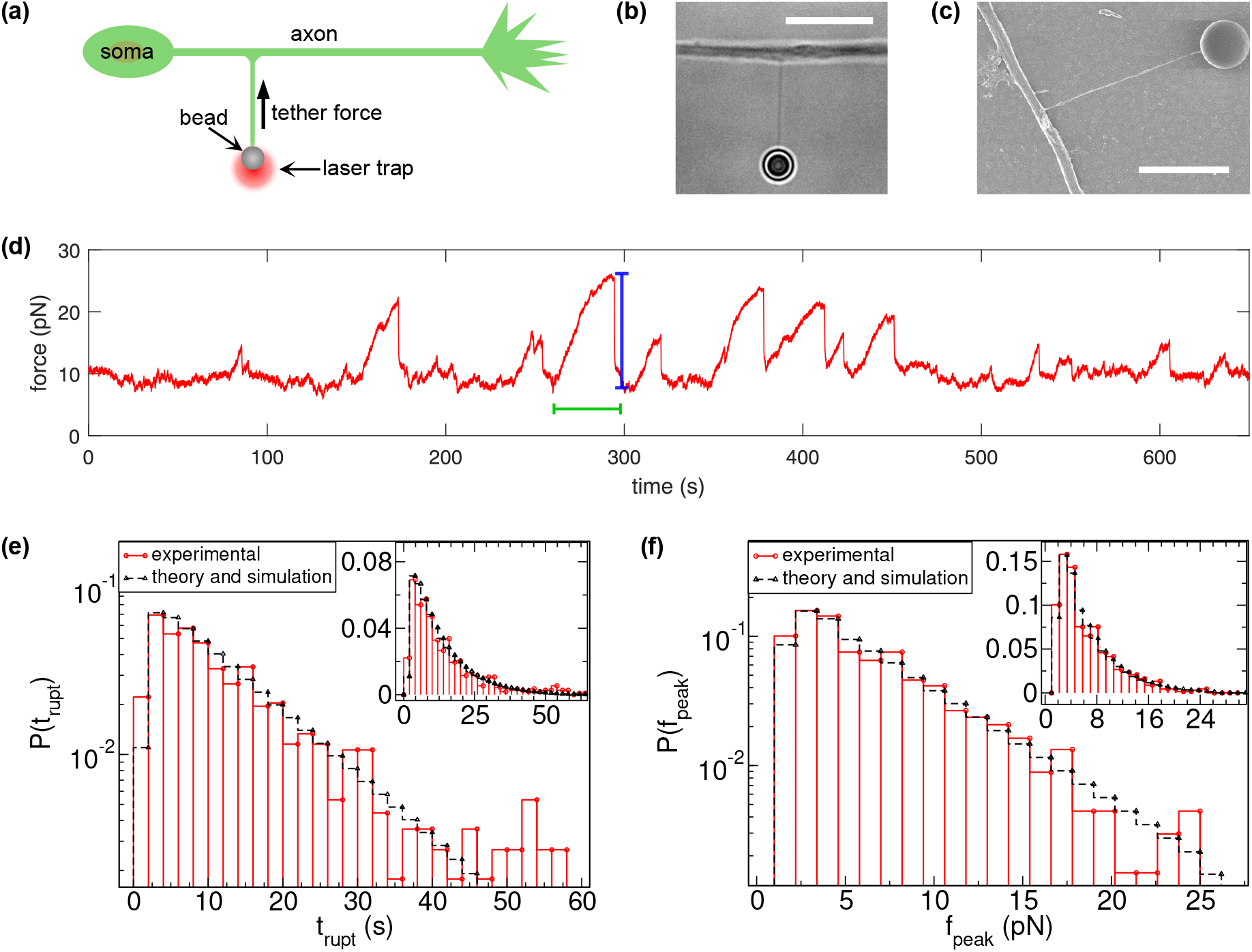
(a) Schematic of the tether pulling experiment. After extracting a tether the distance between the axonal shaft and the optical trap is kept fixed and the force on the trapped bead is monitored. The arrow indicate the positive direction of force. (b) A phase-contrast microscopy image of the trapped bead and the tether (scale bar = 10 µm). (c) A scanning electron microscopy image of an extracted tether (scale bar = 5 µm). This tether had a diameter of 83 nm. (d) Time-series showing the load-fail behavior of a nanotube pulled out of an axon using optical tweezers. (e, f) Histograms (semi-log Y) for the rupture time (*t*_rupt_) as defined by the green bar in (d) and the peak height (*f*_peak_) as defined by the blue bar in (d) obtained using time-series recorded from several axons (34 axons, 80 tethers, 564 peaks). The insets show the data in linear scale. Histograms obtained from stochastic numerical simulations of the theoretical model - explained in the model section - are also shown. The theoretical distributions correspond to the rate of cofilin kinetics (*k*_b_ = *k*_on_ + *k*_off_ = 2*k*_on_, see Eq. 1), with *k*_b_ = 0.25 s^−1^. We consider the actin attachment rate (*k*_a_) and rupture rate (*k*_r0_) to be equal to the rate of cofilin kinetics (*k*_a_ = *k*_r0_ = *k*_b_). Initially, multiple filaments can attach simultaneously with probabilities *p*_1_ = 3*/*7, *p*_2_ = 3*/*7, and *p*_3_ = 1*/*7, where the total number of filaments is *n*_fil_ = 3.

**Figure 2.**
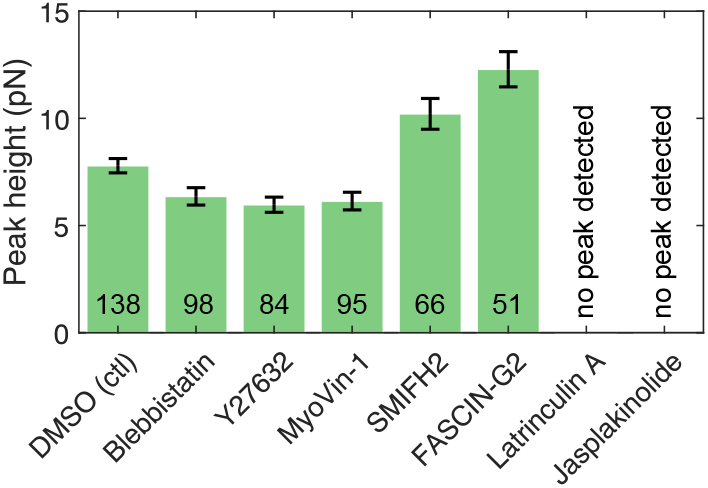
Bar plots showing the average peak height obtained from the distributions of force (maximum force above baseline) when the neurons were exposed to different drugs. The DMSO data is the vehicle control (normal cells). The data for each drug was obtained using multiple axons. The number inside each bin indicates total number of peaks analyzed and the error bars are standard error of the mean.

### ACTIN FILAMENTS SHOW GROWTH AND RETRACTION CYCLES

To explore the role of actin filaments further, we performed fluorescence imaging using cells expressing either LifeAct-mCherry or pCAG-Actin-GFP. As can be seen from the sequence of images and the kymograph shown in Fig. 3a, actin fluorescence exhibit a cyclic growth and decay dynamics within the nanotube (Suppl. Video SV1). As can be seen from the kymograph, the retraction of actin fluorescence in the tube occurs with a maximum velocity near the tip and the velocity decreases towards the base. A quantification of the time evolution of the fluorescence intensity within the nanotube is shown in Fig. 3b. These data show the following. (i) During the “growth phase”, when the fluorescent region grows towards the tip, the intensity at any time point decays monotonically from the base towards the tip (time point i in Fig. 3b). (ii) During the “retraction phase”, regions of localized actin build up are often observed (time point ii in Fig. 3b). These peaks of fluorescence intensity can occur anywhere along the tether and they drift towards the base of the tube. (iii) For the case shown in Fig. 3b, there is hardly any detectable intensity between the retraction front and the tip of the membrane tube (time point ii). This, along with the occurrence of localized fluorescence peaks, suggests that actin retracts as polymerized filaments before undergoing any subsequent depolymerization and regrowth. (iv) After the retraction phase, the intensity becomes more diffused and eventually regains the monotonically decreasing pattern (time point iii in Fig. 3b), and the cycles repeat.

**Figure 3.**
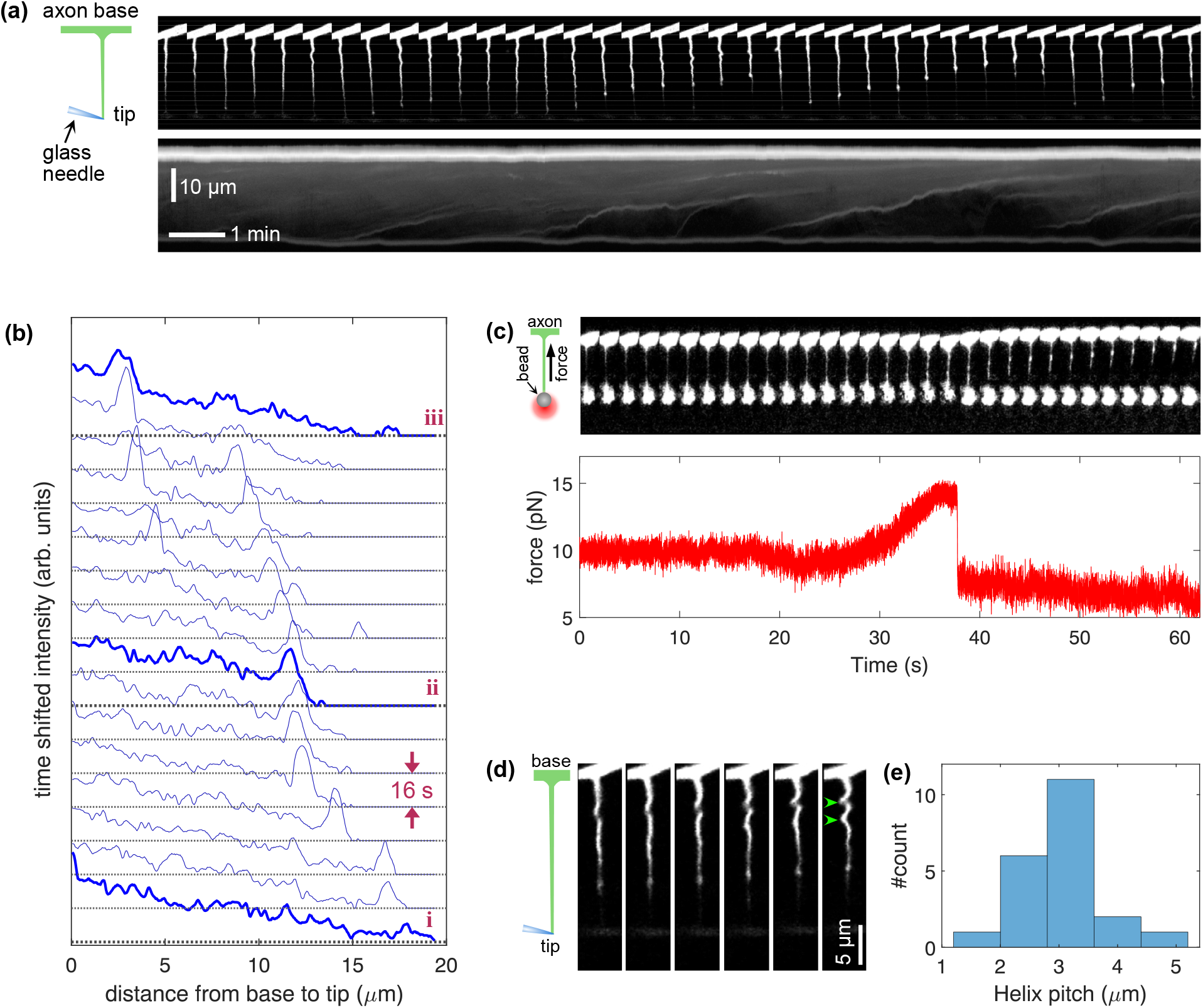
(a) (above) A collage of a sequence of images showing the growth-retraction cycles of fluorescent actin filaments labelled with pCAG-mCherry inside a nanotube. In this case, the tube was pulled out using a glass micro-needle (schematic to the left) and imaged using a confocal microscope. The base of the axon is on top and the tip of the needle is at the bottom (see schematic on the left). (below) Kymograph of the fluorescence intensity for the nanotube shown above. Note that the retraction velocity is a maximum near the tip of the tube and decreases towards the base. The vertical scale bar is 10 µm and horizontal scale bar is 1 min. (b) Sequence of plots of the radially integrated fluorescence intensity along the tether. The lower most curve is the earliest time point, time between curves is 16 s, and a few selected time points are plotted in brighter blue and numbered for clarity. When the actin intensity spans the entire tube, the intensity decreases monotonically from the base, which is to the left, towards the tip of the tube (time point i). As actin intensity recedes, intensity peaks develop and there is hardly any fluorescence at the far end of the tether (time point ii). This suggest retraction of intact filaments as opposed to depolymerisation. The continuous, decreasing profile is recovered once filaments grow back (time point iii) and the cycle repeats. (c) (above) Actin fluorescence in a nanotube extracted using optical tweezers (the trapped bead is towards the bottom). (below) Force exerted on the bead by the nanotube shown above. It can be seen that the force increases when the actin fluorescence spans the tube and drops suddenly when the intensity along the tube becomes discontinuous. This suggest detachment of filaments from the bead. (d) Sequence of images showing helical buckling of actin filaments seen frequently during the retraction phase. (e) Histogram showing the pitch of the helical buckling. Since only a couple of twists are clearly observable along each tether, we have quantified the spacing as shown by the arrow heads in (d). The data was obtained using 4 axons, 6 separate tethers and 21 buckling events.

Fig. 3c shows the optical tweezers measurements performed on a fluorescently labelled cell. In this case, the force increase occurs when the actin intensity becomes continuous and the sudden drop in force coincides with a discontinuity in actin fluorescence. Occasionally, we observe nanotubes where no clear break in fluorescence intensity is seen when the force drops. We presume that in such cases there is still a loss of mechanical continuity due to yielding of filaments within the nanotube. Nanotubes frequently exhibit helical buckling as the actin detach and retracts within the tube (Fig. 3d, and Suppl. Video SV1). The helical buckling as well as the actin growth-retraction cycles can be seen even in extremely long nanotubes as well (*>*50 *µm*) and the wavelength of buckling is similar to that seen in shorter tubes (see Suppl. Video SV2).

Since blocking of myosin II or myosin V activity did not prevent the load-fail behavior, we considered the possibility of actin filaments getting pulled by kinesin or dynein motors on axonal microtubules. Disrupting axonal microtubules by exposure to 10 µM Nocodazole for *>*15 min did not have any noticable effect on the force behavior. Since such a treatment causes only partial disruption of microtubules in axons [14], we also performed experiments using chick primary fibroblasts transfected with tubulin-GFP. Here we could confirm complete dissociation of microtubules and such fibroblast cells, as well as untreated fibroblasts, exhibited active force dynamics similar to that seen in axons (Suppl. Mat. Fig. S5 and S6). As will be discussed in more detail later, our experiments suggest that molecular motors are unlikely to be the generators of force within the actin filaments that invade the nanotubes.

### ROLE OF ADF/COFILIN IN ACTIN-MEDIATED FORCE CYCLES

Since actin stabilization by Jasplakinolide inhibits the force cycles, we investigated the possible role of the actin severing proteins that belong to the ADF/cofilin family [15–19]. Chick DRG neurons contain the ADF (Actin Depolymerising Factor) analogue of cofilin [20]. Imaging using a RFP tag to ADF (using pCAG-ADF-RFP) and GFP tag to actin (pCAG-Actin-GFP) reveal a co-localization and highly correlated dynamics of ADF and actin within nanotubes (Fig. 4a,b and Suppl. Video SV3). Next, we expressed either a constitutively inactive form of ADF (ADF S3E) [21] or a constitutively active form (ADF S3A) [22], and recorded the force time-series. As shown in Fig. 4c, competition with inactive ADF decreased the peak height while the peak duration is increased (Welch’s t-test was performed for statistical significance and significance was assessed at *α* = 0.01). Remarkably, in some cases the peak duration lasted for as much as 250 s with the force remaining close to a saturation value (suppl. mat. Fig. S7). Such long rupture times with the force remaining at saturation level were never seen in normal cells. Expression of the active form of ADF, on the other hand, did not show any significant difference in peak height or peak duration compared to control (see Materials and Methods for statistical analysis).

**Figure 4.**
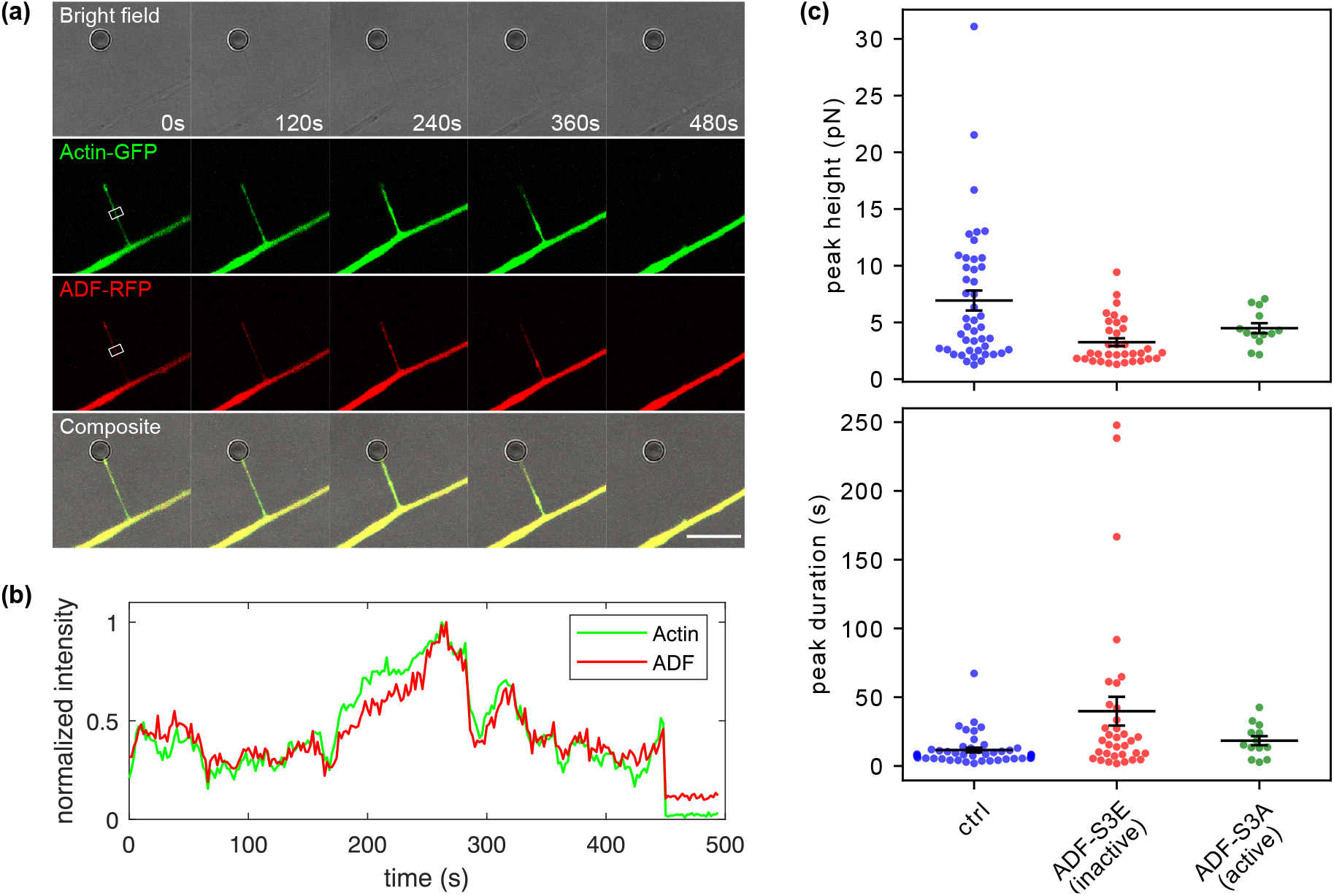
(a) Bright field and fluorescence snapshots of an axonal tether pulled from a neuron co-transfected with pCAG-Actin-GFP (for actin) and pCAG-ADF-RFP (for ADF). The composite image shows the extent of co-localization of ADF with actin in the tether (Suppl. Video SV3). The scale bar is 10 µm. (b) Plots of the time evolution of the normalized intensity for actin and ADF calculated within the white boxes shown in (a) for each channel. (c) Beeswarm plot of peak height and peak duration when neurons were transfected with a constitutively inactive form of ADF (ADF-S3E) and constitutively active form of ADF (ADF-S3A), with soluble GFP (pCAG-GFP) used as marker for successful transfection. Horizontal bars are mean values and error bars are standard error of means of respective distributions. We observed a significant decrease in peak height and significant increase in the peak duration, compared to control, with ADF-S3E. No significant change was observed for cells trasfected with ADF-S3A (Welch’s t-test: *α* = 0.01, refer to Materials and Methods for details of statistical analysis).

### A STOCHASTIC MODEL WITH FILAMENT SHRINKAGE REPRODUCES THE LOAD-AND-FAIL BEHAVIOR

We propose to analyze the statistics of the load and fail cycles of the nanotube based on a stochastic model involving actin filaments attaching to and detaching from the nanotube tip, and progressive filament shrinkage due to the binding of cytoplasmic actin associated proteins, thereafter labeled ADF/cofilin for conciseness, see Fig. 5 (a) for a sketch of the model. The assumption of protein-induced filament shrinkage can be thought of as a generic mechanism for the generation of pulling forces. We show in the Suppl. Mat. that the induction of torsional and/or curvature stress which can be inferred from the helical shape of detached filament (Fig. 3d) can, in the linear regime, be reproduced by an effective shrinkage for attached filaments. The growth of actin filaments is not included in the model to limit the number of parameters, and therefore, the model does not capture the time delay between successive loading events.

**Figure 5.**
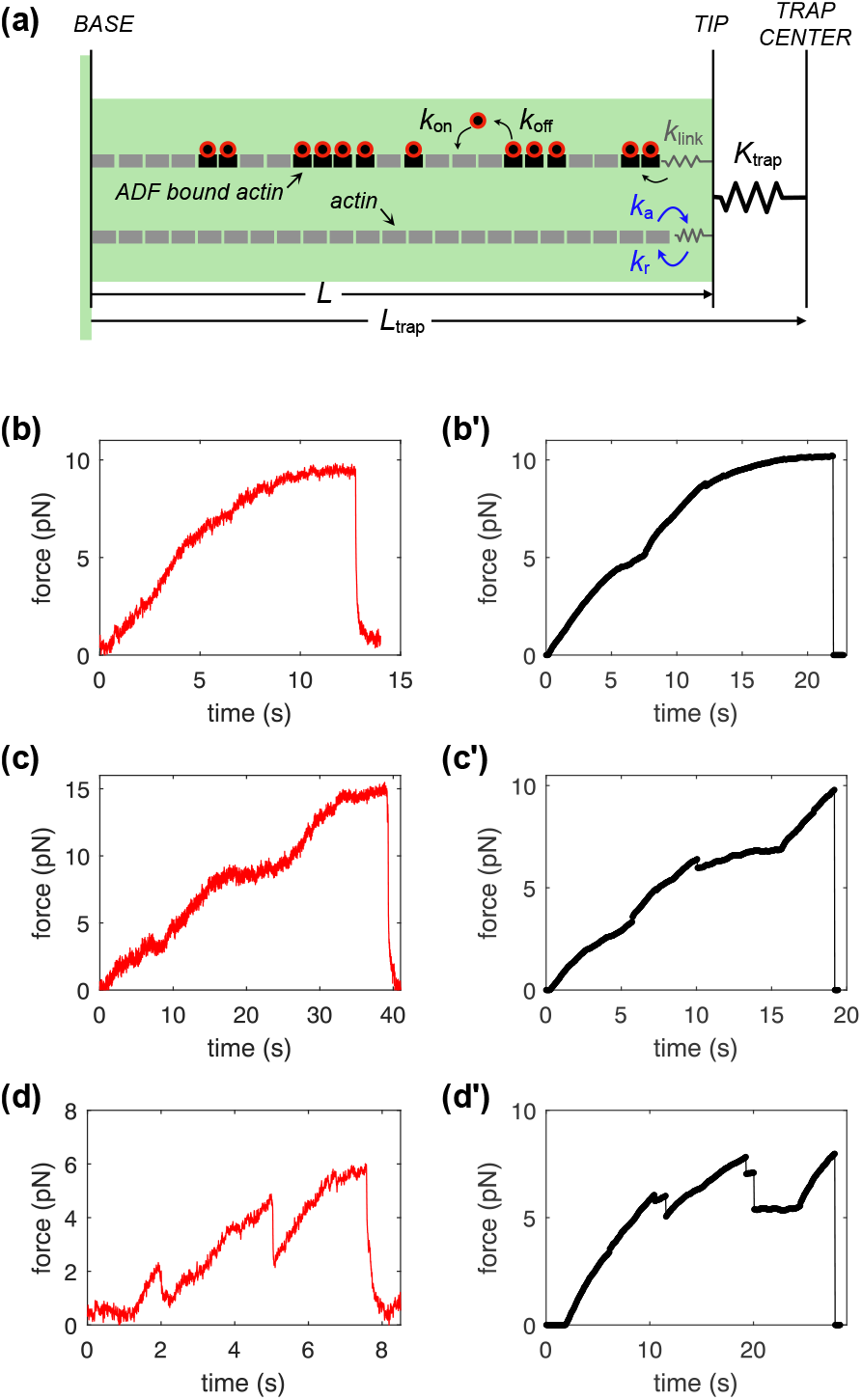
(a) Sketch of the theoretical model. The membrane tether is shown in light green. The optical trap is modeled as a spring of stiffness *K*trap. Actin filaments (gray dotted lines) emerging from the base of the tether can attach to and detach from the tether tip with rates *k*_a_ and *k*_r_, respectively. The linker between the filament and the tip is a spring of stiffness *k*_link_. ADF/cofilin (red circles) can bind to a naked monomer (gray rectangle) with a rate *k*_on_ and unbind from the decorated monomer (black rectangle) with a rate *k*_off_. ADF/cofilin-decorated monomers have a smaller length than ADF/cofilin-free monomers. The net tether force is obtained by multiplying the distance between the tip and the center of the trap by the trap stiffness *K*_trap_. Lower panels: Qualitative comparison between particular features of the variation of the tether force with time seen in experiments (b, c, d) and in the theoretical model (b′, c′, d′). (b, b′) show force saturation, (c, c′) show changes of slope of the force with time and (d, d′) show partial force drops requiring the presence of multiple filaments. For the theoretical force-time profiles, we consider *k*_a_ = *k*_r_ = *k*_b_ and *k*_b_ = 0.25 *s*^−1^.

- **Shrinkage of individual filaments:** As the force steadily increases with time between failure events, ADF/cofilin binding can be thought of as an almost continuous process. Considering a filament made of *N* actin subunits, each of length 𝓁, the filament length prior to any ADF/cofilin binding is *L*_0_ = *N*𝓁. We assume that a single ADF/cofilin can bind to a single actin monomer [23, 24] and decreases its rest (stress free) length by an amount *δ*𝓁. The rest length of the filament is thus *L* = *L*_0_(1− *ϵδ*𝓁*/*𝓁), where *ϵ* is the faction of subunits carrying ADF/cofilin. The time evolution of *ϵ* satisfies the kinetic equation 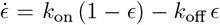, where the ADF/cofilin binding and unbinding rates *k*_on_ and *k*_off_ could depend on local properties such as the mechanical stress. For simplicity, we assume here that they do not, so that starting with a ADF/cofilin-free filament at *t* = 0, we have

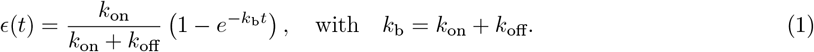
- **Force exerted by an individual filament:** An empty membrane tether held by an optical trap exerts a force 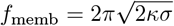on the trap, where *σ* is the membrane tension and *κ* is the membrane bending rigidity [25]. The length of such tube is then *L*_memb_ = *L*_trap_− *f*_memb_*/K*_trap_, where *K*_trap_ is the stiffness of the trap and *L*_trap_ is the (imposed) distance between the center of the trap and the base of the tether. An actin filament that reaches the tip of the tether and attaches to it has an initial length *L*_0_ = *L*_trap_. As this filament shrinks, the trap force increases: *F* = *f*_memb_ + *f*_a_, where the contribution of actin to the force is *f*_a_(*t*) = *k*_eff_*L*_0_*ϵ*(*t*)*δ*𝓁*/*𝓁, where *k*_eff_ = 1*/*(1*/K*_trap_ + 1*/k*_link_) is an effective stiffness that also account for the stiffness *k*_link_ of the actin filament and the linker between the filament and the tether tip. In the limit *k*_link_ ≫ *K*_trap_, which we adopt here, we have *k*_eff_ ≃ *K*_trap_. Finally, the evolution of the force generated by a single filament is:

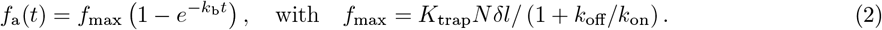 This force will increase until it either saturates and eventually drop when the filament detaches or drops before saturation. By choosing a particular rupture kinetics, one may derive the distribution of rupture forces and rupture times as is measured experimentally. As shown in the Suppl. Mat., these distributions can be computed analytically for constant rupture rate, or for mechano-sensitive rupture following Bell’s statistics (rate increasing exponentially with the force). Quantitative analysis (detailed in the Suppl. Mat.), shows that the single filament picture – even with mechano-sensitive rupture – is unable to reproduce the salient features of the force and time probability distributions shown in Fig. 1e,f, namely the presence of a peak at small force (and time), and an exponential distribution for large force (and time). Indeed, the single filament picture rather predicts an exponential distribution (for constant unbinding rate) or a peak with short range decay (for mechano-sensitive unbinding). Furthermore, some peculiar features of the force trace, in particular the existence of partial rupture events after which the force remains above the baseline (see Fig. 5 (d, d’)) suggests that multiple filaments are involved in the force generation process.
- **Force exerted by multiple filaments:** The situation where several filaments can be attach to the tether tip simultaneously involves non trivial collective effects and must be handled numerically. Indeed, filaments binding at different times shrink at different rates. Filaments shrinking the slowest will be compressed by the fast shrinking filaments, affecting the net tether force in complex ways. To model this, we assume that a given filament *i* attaches under no tension at time *t*_*i*_ with a length equal to the tether length at the time of attachment *L*(*t*_*i*_). As ADF/cofilin binds to this filament, its rest length *L*_*i*_(*t*) decreases, and it pulls on the tip with a force *f*_*i*_(*t*) = *k*_*i*_(*L*(*t*) − *L*_*i*_(*t*)), where *k*_*i*_ accounts for the stiffness of the filament and the linker protein connecting the he linker protein connecting the filament to the tip. The tether force is then *F* (*t*) = *f*_memb_ + ∑_*i*_ *f*_*i*_(*t*), which can be written as:

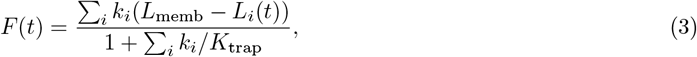

and the force on the *i*th filament is expressed as

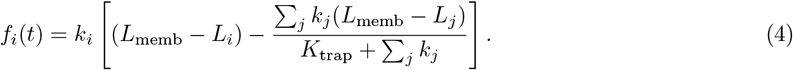 We note that if all filaments and linkers are identical (*k*_*i*_ = *k*) and bind simultaneously, the total average force is *F* = *n*_fil_ *f*_1_ = *n*_fil_ *k*(*L*_memb_ − *L*_1_)*/*(1 + *n*_fil_ *k/K*_trap_), where *n*_fil_ is the total number of filaments. In the limit *n*_fil_*k* ≫ *K*_trap_, the total average force becomes *F* = *K*_trap_(*L*_memb_− *L*_1_), which is independent of the number of filaments *n*_fil_. On the other hand, soft filaments (*n*_fil_*k* ≪ *K*_trap_) produce a force (*F* = *n*_fil_ *k*(*L*_memb_ − *L*_1_)) proportional to the number of filament in this situation.
- **Numerical simulations:** We use the Gillespie algorithm [26] to model the system. This algorithm considers agent based processes that incorporates stochasticity in the individual level for both the binding and unbinding of ADF/cofilins and the attachment and detachment of the filaments on the tether tip. We consider each event as a reaction that occurs in real time based on its respective rate. Let us consider a total number of filaments *n*_fil_, with *n*_a_ attached filaments and *n*_d_ = *n*_fil_ − *n*_a_ detached filaments at a given time *t*. We first calculate the total propensity of all reactions, which can be written as

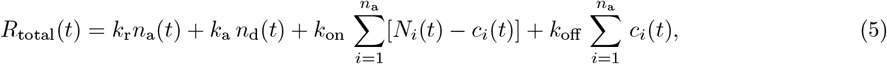

where *c*_*i*_ and *N*_*i*_ are the number of bound ADF/cofilins and the total number of available sites on the *i*th filament, respectively. The rupture and attachment rates of a filament are denoted by *k*_r_ and *k*_a_, respectively. Once a reaction occurs, the next reaction - whichever it is - will occur Δ*t* time later which is calculated as Δ*t* = log(1*/r*)*/R*_total_ where *r* ∈ [0, 1) is a random number chosen from a uniform distribution. A reaction occurs with a probability determined by the ratio of the reaction rate to the total rate. For example, the detachment of the *i*th filament occurs with a probability of *k*_r_*/R*_total_. Each force evolution with time (until rupture) is considered as an ensemble in our statistical analysis.

The results of the simulation for single filaments with mechanosensitive rupture (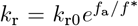) are shown are shown in the Suppl. Mat. It compares very well with the analytical solution also discussed in the Suppl. Mat.. However, as discussed earlier, the single filament case with the mechanosensitive rupture is not sufficient to fit the experimental data (see Suppl. Mat. Figs. S10-S11). We found that allowing three or four filaments tobe bound simultaneously was appropriate and we adopt *n*_*fil*_ = 3 here. We call the probability that a number *i* ∈ [1, 3] filaments are bound at the start o(f an) force generation event as *p*_*i*_. If the filaments are distinguishable, *i* filaments can attach simultaneously in 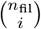 ways. With *n*_fil_ = 3, there are 7 possible configurations, with initial probabilities *p*_1_ = 3*/*7, *p*_2_ = 3*/*7, and *p*_3_ = 1*/*7. We adopt these initial probabilities here for simplicity, but it should be noted that the distributions are not very sensitive to the chosen values of *p*_*i*_. For instance, we demonstrate how setting the initial probability to 1 for one, two, or all three filaments affects both the distributions shown in Fig. S 16. The simulation results for the peak force and rupture time distributions show a good match with the experimental data (see Fig. 1 (e,f)). Moreover, as shown in Fig. (5 (b-d), the broad variety of features of the load and fail behavior seen experimentally are well reproduced by the simulations. An example time series and a more detailed comparison of these are shown in Figs. S 12-14.

## DISCUSSION

### Comparison with filopodia contraction

Although membrane nanotubes pulled out of axons resembles filopodia in some ways, the contractile mechanism we observe is distinct from the several mechanisms proposed earlier for filopodial contractility, as summarized below. Based on traction force microscopy, Chan *et al*. demonstrated that filopodia exhibit repeated load and fail cycles [9]. They modelled mechanosensitive contractility by invoking myosin II mediated pulling at the base of the filopodia, although the involvement of myosin II in force generation was not explored in their experiments. In general, the role of myosin II in filopodial contraction is not clear with different studies on different cell types yielding different results [10, 27–29]. In our case, multiple drug experiments clearly rule out any role for myosin II in the contraction of extracted nanotubes. Another mechanism proposed by Bornschlögl *et al*. for pulling force generation in filopodia invokes frictional coupling between the retrograde flow of actin filaments in the cortex and actin filaments extending from the filopodium [11]. Such a mechanism where actin filaments in the tube are pulled at the base due to cortical retrograde flow can be ruled out in our case because the flow velocity is nearly zero at the base of the nanotubes (see kymograph of Fig.3a). Unconventional myosins like myosins V & VI have also been tested in the case of retracting phagocytotic filopodia, and knock-out of either of these did not affect retraction, and no force measurement is reported [30]. Based on optical tweezers measurements, Farrell *et al*. have proposed a force generation mechanism that is mediated by end-tracking linkages between the actin filaments and the bead [10]. Generally, such linkages occur via formins, and formins can also act as force generators [31]. Such mechanisms are unlikely in our case as inhibition of formins did not abolish the force events (Fig. 2). More recently, based on the rotation of actin filaments inside filopodia, knock-down experiments, and theoretical modelling, Leijnse et al. have argued that unconventional myosins like myosin V & X may be involved in the helical buckling of the actin bundles [12]. While such active processes can account for contractile stresses, how the cyclic growth and retraction process is regulated within the membrane tube is not fully understood. Several studies performed on filopodia have highlighted the role of actin severing proteins belonging to the ADF/cofilin family in regulating actin dynamics [18, 32, 33]. Our experiments using mutant ADF constructs suggest that stresses are generated by ADF/cofilin binding leading to contractile forces in actin bundles, subsequent disruption of filaments, followed by repolymerization. This process is detailed below.

### ADF/cofilin based contractility hypothesis

We observe that ADF co-localizes with the dynamic actin within the nanotubes (Fig. 4a,b). Reduction in ADF activity caused by competition with an inactive form which competes with native ADF results in very long lived force events and reduced peak force when compared to control cells, suggesting a role for ADF in force generation (Fig. 4c). No significant changes to the load-fail response is seen when a constitutively active form of ADF is over-expressed (Fig. 4c). This can be understood if the concentration of the native form of acive ADF is already high enough to saturate the filaments.

The ADF/cofilin based mechanism proposed here can account for the sustained cyclic load-fail behavior seen in the nanotubes as follows. ADF/cofilin is known to decorate actin filaments in filopodia and drive disassembly [18, 32, 33]. Freshly formed actin filaments are composed of ATP-bound monomers [34] and are prone to ADF/cofilin-induced severing only after they have been converted to the ADP-bound form [35]. This causes a delay between the initial growth of actin within the tube, subsequent stress generation due to ADF/cofilin binding, and the eventual severing and disintegration of filaments. The G-actin thus generated undergoes exchange of nucleotide to participate in fresh filament formation, completing the cycle. The time taken for fresh actin to polymerize, hydrolyse ATP, and bind cofilin, gives rise to the “dead-time” between successive force events.

Based on the above observations, We hypothesize that stresses generated by ADF on actin filaments can also account for the measured contractile forces. It is known that the binding of ADF/cofilin results in the generation of torsional stress that eventually causes severing [19, 36–39]. The torsional stress manifest as an over-twisting of actin filaments leading to a ≈ 25% reduction in the actin pitch [33, 37, 38]. We argue that this over-twisting of actin filaments is accompanied by a normal contractile stress resulting in a reduction in the length of each actin filament.

A correlation between longitudinal stress in actin filaments and cofilin-induced severing was demonstrated using an *in-vitro* system [40]. There, an externally imposed longitudinal tension on actin filaments held using optical tweezers markedly decreased the severing probability of cofilin. As a corollary, in the mechanism we propose here, the binding of ADF on actin can induce conformational changes leading to generation of tension when the ends are constrained between boundaries. Such effects, are expected due to allosteric reasons, as was predicted based on the experimentally observed plasticity of actin filaments [16, 41]. Remarkably, an allosteric force generation mechanism has been reported recently to explain the generation of longitudinal tension in microtubules held in a double optical trap when the microtubule associated protein tau was allowed to bind [42].

### Simple model of ADF/cofilin-mediated contractility

Here, we propose a simple model where ADF/cofilin binding imposes actin filaments to shrink, generating a pulling force on the tether tip. In this model, actin filaments are assumed to be straight, but in practice, this effect can also be obtained if ADF/cofilin binding imposes a helical twist to the filament (see Suppl. Mat.). Owing to the short-time linear increase of tether force with time seen in our experiments, we assume that this shrinkage is constant per bound protein, and that binding follows a first order kinetics that eventually saturates to a value determined by the ratio of the protein binding to unbinding rates.

Our stochastic model includes ADF/cofilin binding on actin, which generates contraction, and actin attachment and detachment to the tip of the membrane tube. We show that qualitative features of the time dependent force (such as abrupt partial drops of force that do not reach the baseline, Fig. 5 d,d’ and Fig. S 12) requires that several (at least three) actin filaments can be bound simultaneously to the tether tip. With this assumption, the model is able to quantitatively reproduce the distribution of peak force and rupture time measured experimentally. Moreover, certain features of the temporal variation of the force, such as sharp increase and decrease in slope, are also reproduced (Fig. 5 c,c’ and Fig. S 13).

This simple model does not account for the time delay between contraction events, which depend on actin polymerisation and associated ATP to ADP conversion, but concentrates on the statistics of the events themselves. Furthermore, the model does not include tension-dependent ADF/cofilin binding/unbinding or actin attachment/detachment. While these features are not necessary to reproduced the peak force and duration statistics, they might nevertheless be important for other features, such as the anticorrelation between peak force and duration when cells are transfected with a constitutively inactive form of ADF (Fig. 4c).

### Conclusion

In summary, we demonstrate myosin-II independent and actin-based cyclic contractility in nanotubes pulled out of axons. Axons form a convenient system for quantitative force analysis as they lack any significant motility that can corrupt the force data. This allows us to make comparisons between the statistical properties of experimentally and theoretically obtained load and fail responses. The ADF/cofilin based mechanisms we have proposed here can be further tested using *in-vitro* systems using optical tweezers. To what extent such mechanisms are involved in actual filopodial dynamics is also an area for further exploration.

## MATERIALS AND METHODS

### Preparing cell culture media

L-15 medium (Gibco 21083-027) thickened with 6 mg/ml Methyl Cellulose (Colorcon, ID34516) and supplemented with 10% (v/v) Fetal bovine serum (Gibco, 10100-147), 6 mg/mL D-Glucose (Sigma G6152), 20 ng/ml Nerve Growth Factor (NGF-7s, 13290-010, Thermo Fisher Sci.) and 10 µl/ml Penicillin-Streptomycin-Glutamine (Gibco 10378-016).

### Neuronal cell culture

Chicken eggs were incubated for 8-9 days at 37 ^*°*^C, then the embryo from the egg was dissected in HBSS buffer with Ca^++^/Mg^++^ (Gibco, 14025-092) for isolation of Dorsal Root Ganglia (DRG). The extracted DRGs were incubated in 0.25 % Trypsin-EDTA (Gibco, 15400-054) at 37 ^*°*^C for 20 min. The cells were isolated using centrifugation and the dissociated cells were re-suspended in 100 µL OPTI-MEM 1X solution (Gibco, 31985-062) by gentle pipetting. The suspended cells were then plated on glass cover-slips attached to punched 35 mm petri dishes with 2 mL of pre-warmed cell culture media and incubated at 37 ^*°*^C for 24-48 hrs. The glass cover slips were thoroughly cleaned by sonication in a detergent solution (Extran, MA03 Merck, Millipore), rinsed several times with Millipore deionized water, dried, sterilized under UV and used without any further treatment. The cell culture medium was then replaced with culture medium lacking Methyl Cellulose 30 min before the start of the experiments.

Electroporation method was used for transfecting the cells with external plasmids or siRNA. To do the transfection with plasmid, before culturing the cells 10 µg of plasmid/s of interest is added to the dissociated cell suspension (in 100 µL OPTI-MEM) and electroporated using super electroporator (NEPA GENE, NEPA21 Type II). For siRNA (ADF-1 - 5’-GUGGAAGAAGGCAAAGAGAUU-3’, Lac Z - 5’-CGUCGACGGAAUACUUCGAUU-3’, Integrated DNA Technology) transfection, it was added to dissociated cell suspension in 30 nM concentration before electroporation.

Lifeact-mCherry and pCAG-actin-GFP was used for fluorescene imaging of actin inside membrane nanotubes. Fluorescent mutant of ADF S3A and ADF S3E was not available so it was mixed with empty vector GFP (pCAG-GFP) as a marker for trasfected cells. siRNA-ADF was used to inhibit ADF production in the cell and siRNA-LacZ (scrambled RNA) was used as control. GFP-tubulin was used for tagging microtubule.

### Pulling membrane nano-tubes from axons

A home built optical tweezers was used for nanotube pulling and force measurement experiments. The setup consisted of a 1W CW DPSS Nd:YAG Laser (Laser Quantum, Ventus 1064+MPC6000) and an Olympus IX71 microscope with a UplanFL N 100X/1.3 Oil objective and custom fitted with IR-reflecting dichroics - one to reflect the laser light into the objective and another to detect the back-scattered laser light via a side port. The microscope was equipped for fluorescence imaging using an EMCCD camera (Andor, iXon DU-885K). Carboxylated polystyrene beads (Spherotech, 2.8 µm) were trapped and used for pulling membrane nanotubes. A motorized XY positioning stage (PRIOR Scientific, H117) was used to displace the sample with respect to the fixed laser trap. The sample chamber was maintained at 37^*°*^C using a home-developed electric incubator. Displacement of a trapped bead from the fixed center of the trap was measured by imaging the back-scattered light on to a Quadrant Photodiode coupled to pre-amplifiers (OSI Optoelectronics, Spot-9DMI) and read into a computer using a 14 bit DAQ card (National Instruments, USB-6009). Trap calibration was done using the standard Power Spectrum method at different laser powers. The trap stiffness at the 90 % laser power (used during experiment) was 100 ± 16 and 90± 14 pN/µm along X and Y axes respectively. For some experiments a glass micro-needle with motorized micro manipulator (Sutter Instruments) mounted on a Leica SP8 confoal microscope was used to pull nano-tubes. Alternatively, nano-tubes were extracted using optical tweezers and the bead was made to stick non-specifically to the cover slip by lowering the trap position and then transferred to the confocal microscope for fluorescence imaging of actin dynamics.

### Drug treatments to cells

All the drugs were dissolved in DMSO, and used with following concentrations. 1 µM Latrinculin A (Sigma-Aldrich, L5163) to depolymerize f-actin, 10 µM Jasplakinolide (Invitrogen, J7473) to stabilize f-actin, 20 µM Blebbistatin (Sigma-Aldrich, B0560) to inhibit myosin II activity, 50 µM MyoVin-1 (CALBIOCHEM, 475984) to inhibit myosin V activity, 40 µM FASCIN-G2 (Xcess Biosciences, M60269-2S) to inhibit actin cross-linking, 20 µM SMIFH2 (Sigma-Aldrich, S4826) to inhibit formin activity, 50 µM CK-666 (Sigma-Aldrich, SML0006) to inhibit Arp 2/3 activity, 16.6 µM Nocodazole (Sigma-Aldrich, M1404) to depolymerize microtubule, 10 µM Y27632 (Sigma-Aldrich, Y0503) as Rho kinase inhibitor, 20 µM ML-7 (sigma-aldrich, I2764) as a myosin light chain kinase (MLCK) inhibitor. All controls were vehicle (DMSO) controls.

ATP depletion was performed by exposing the cells to a medium containing L-15 + 10 mM Sodium azide (Riedel-de Haën 13412) + 10 mM 2-Deoxy-D-glucose (Sigma D6134) for 30 minutes.

### Statistical analysis

As sample sizes were different and variance may not be same, Welch’s t-test was performed for statistical significance. One-tailed Welch’s t-test was performed to check the decrease in peak height and increase in peak duration when cells were transfected with ADF-S3E, the obtained p-values were 0.00013 and 0.0057 respectively. Two-tailed Welch’s t-test was performed to check any change in peak height and peak duration when cells were transfected with ADF-S3A, the obtained p-values were 0.0163 and 0.0831 respectively Statistical significance was assessed at *α* = 0.01.

## ACKNOWLEDGMENT

We acknowledge J R Bamburg for generously sharing all ADF constructs and for helpful suggestions. We are grateful to Jacques Prost for illuminating discussions. We thank Satyajit Mayor for donating Formin and Arp2/3 drugs, as well as protocols. We thank Aurnab Ghose for discussions and for help with ADF cloning work. We Thank K M Yatheendran for performing scanning electron microscopy. The authors acknowledge support through The Wellcome Trust DBT India Alliance (grant IA/TSG/20/1/600137). T.S. and P.S. acknowledge support from the European Research Council (ERC) ERC-SyG (Grant agreement ID: 101071793).

## Notes

### Competing Interest Statement

The authors have declared no competing interest.

