## Supplementary material for "Myosin II independent contraction of actin filaments in membrane nanotubes": Supplemantary Information

### Supplementary material
### Myosin-II independent contraction of actin filaments in membrane nanotubes

Md Arsalan Ashraf[1,†], Tapas Singha[2,†], Susav Pradhan[1], Serene Rose David[1], Pierre Sens[2], and Pramod Pullarkat[1]
[1] *Raman Research Institute, C V Raman Avenue, Bengaluru, 560080, India and*
[2] *Institut Curie, PSL Research University, CNRS UMR 168; F-75005 Paris, France*

#### CAPTIONS FOR VIDEOS

**Supplementary Video 1:** This timelapse video shows the growth and retraction dynamics of fluorescent actin (pCAG-mCherry) filaments within a tether pulled using a glass needle. The imaging was performed using a Leica SP8 confocal system.

**Supplementary Video 2:** Timelapse video showing the actin growth and retraction dynamics within a tether of length greater than 50 µm pulled using a microneedle. The tip of the microneedle, which is coated with FM4-64 fluorescent dye (only faintly visible), is indicated by the arrow. Helical buckling with periodicity similar to that of short tethers can also be seen in such very long tethers.

**Supplementary Video 3:** Timelapse video of simultaneous imaging of fluorescent actin (pCAG-Actin-GFP) and fluorescent pCAG-ADF-RFP. The tether was extracted using optical tweezers and the bead was made to stick to the coverslip to enable imaging using a confocal microscope (Leica SP8). Simultaneous bright field recording performed using the trans-PMT is also shown. The final image (lower-right) shows the merged video for all three channels.

#### I. ADDITIONAL DATA

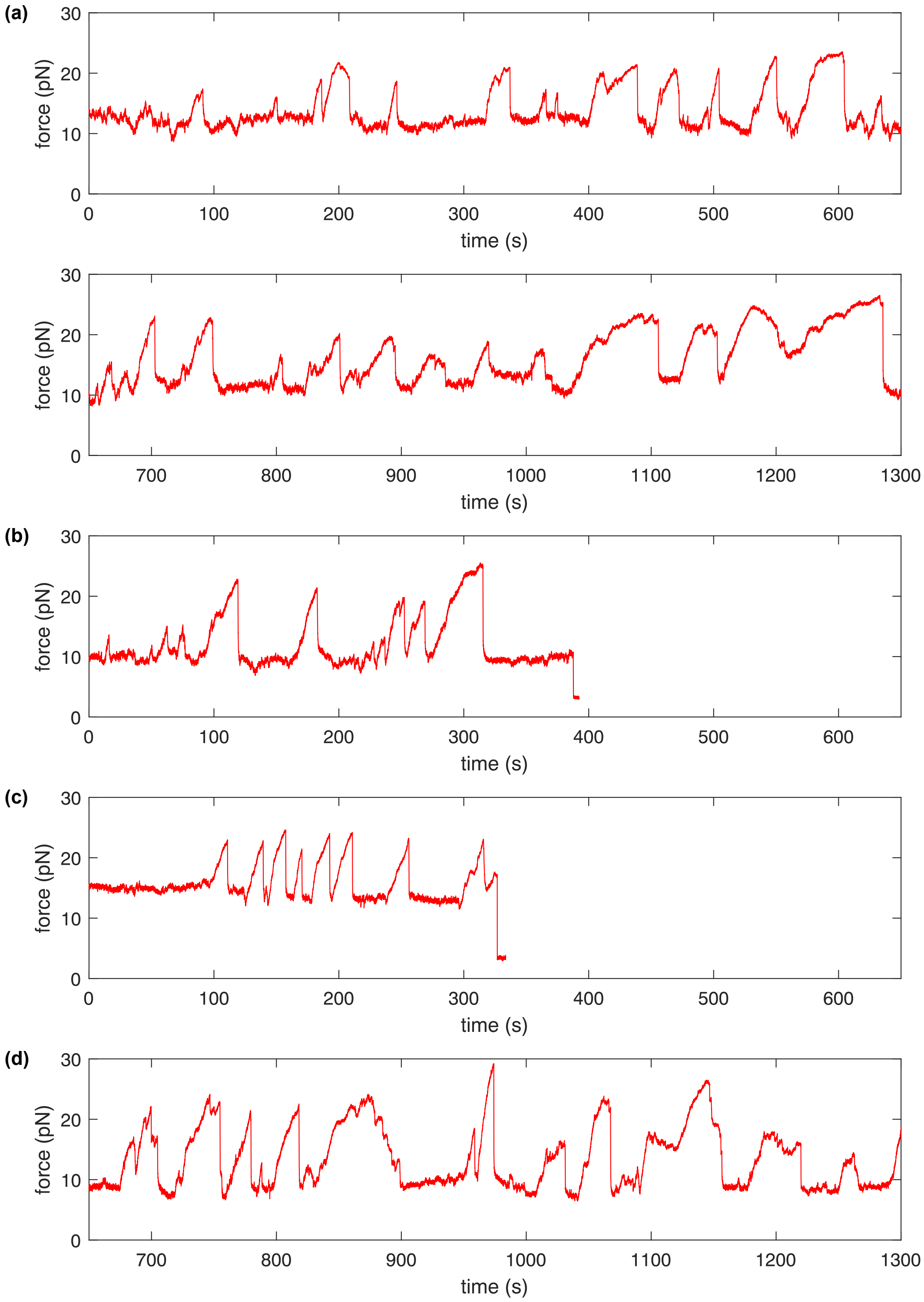


Fig. S 1. Time series data showing tether force from different axons (a,b,c,d). (b)&(c) shows the time series where the tether broke towards the end.

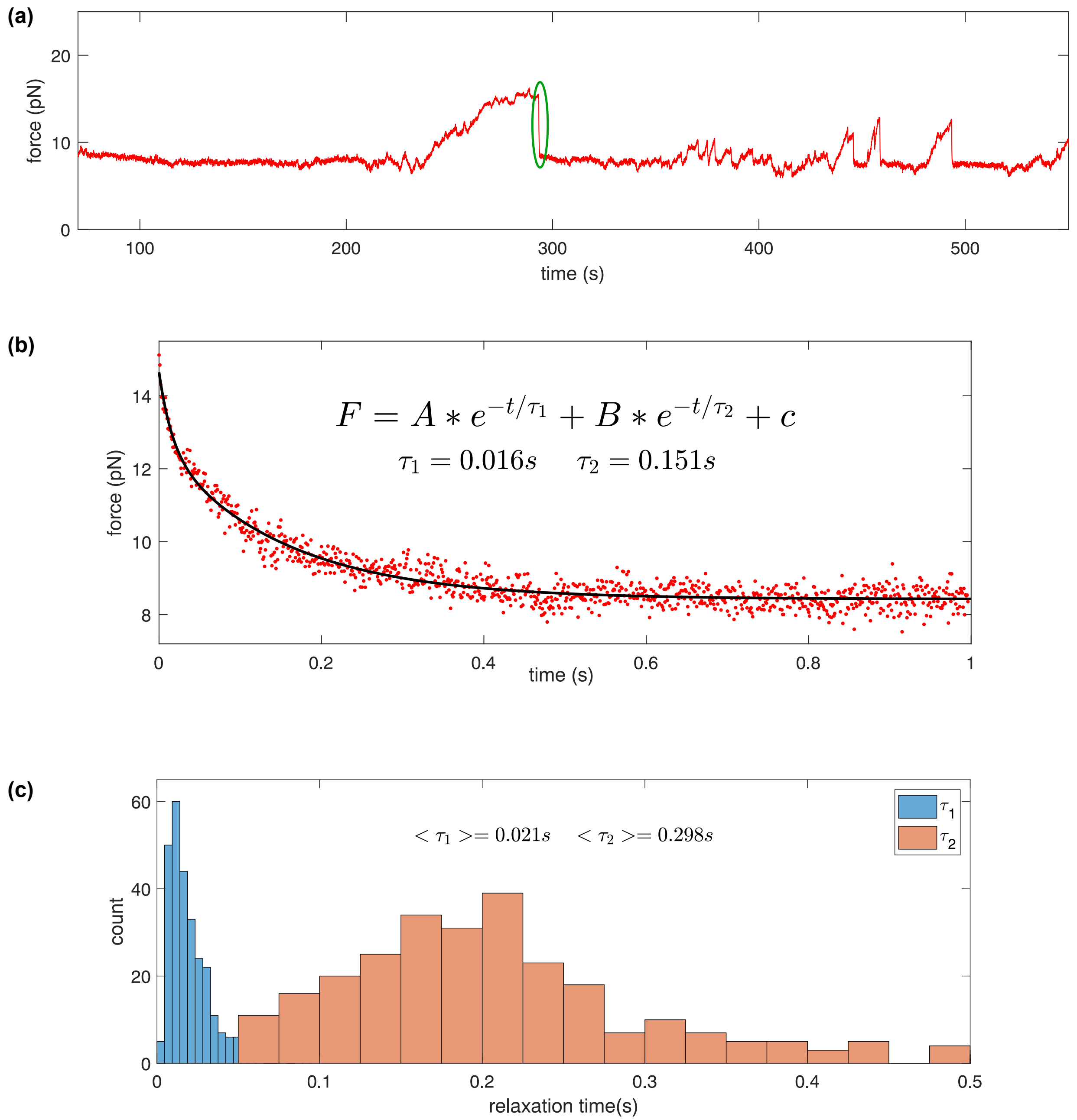


Fig. S 2. (a) Time series data showing active force peaks. One of the force collapse is indicated by green enclosed region. (b) Plot of the force collapse event shown in (a) (red scatter plot) and fitted curve (black line) using the double exponential equation shown in the plot. The two time scales ($\tau_1$ and $\tau_2$) obtained from the fit is also shown. (c) Histogram of $\tau_1$ and $\tau_2$ obtained using 280 force collapse events. $\langle\tau_1\rangle$ and $\langle\tau_2\rangle$, shown as text in the plot, are the mean values of $\tau_1$ and $\tau_2$ respectively. (There are few data points beyond 0.5 s, those are considered as outlier and not shown in the histogram.

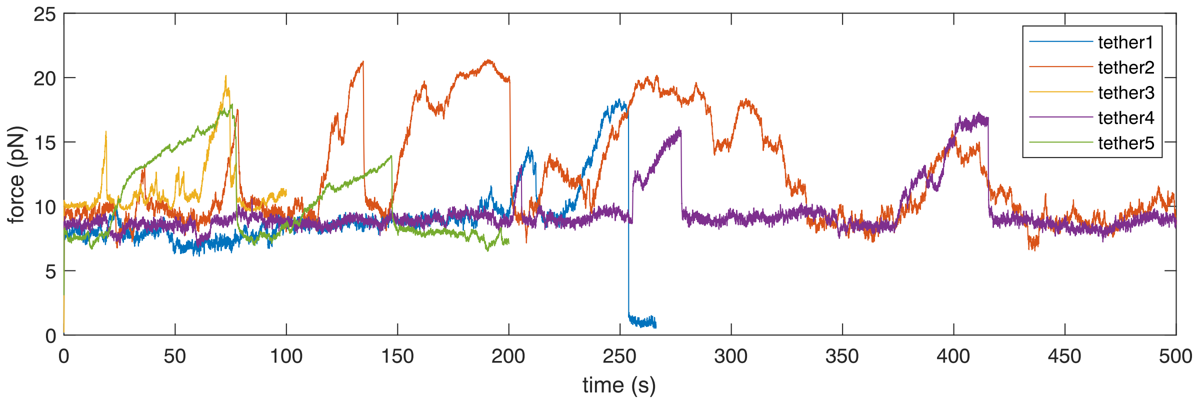


Fig. S 3. Overlayed force-time plots (data from five tethers) to show the occurrence of active peaks in presence of a combination of ML7 (MLCK Inhibitor) and Y27632 (ROCK Inhibitor). ML7 and Y27632 working concentrations were 20 $\mu$m and 10 $\mu$m respectively.

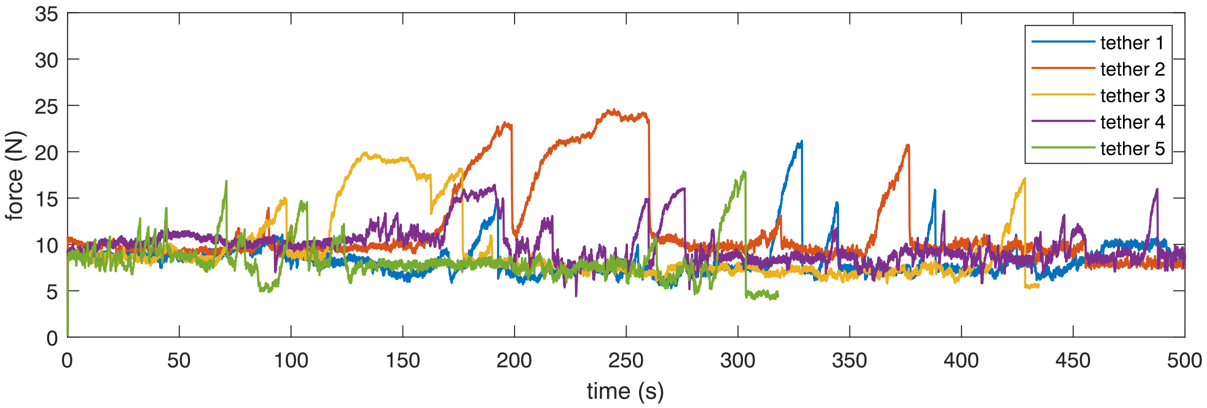


Fig. S 4. Overlayed force-time plots (data from five tethers) to show the occurrence of active peaks in presence of CK666 drug (inhibitor of Arp 2/3 protein), at a concentration of 50 $\mu$M.

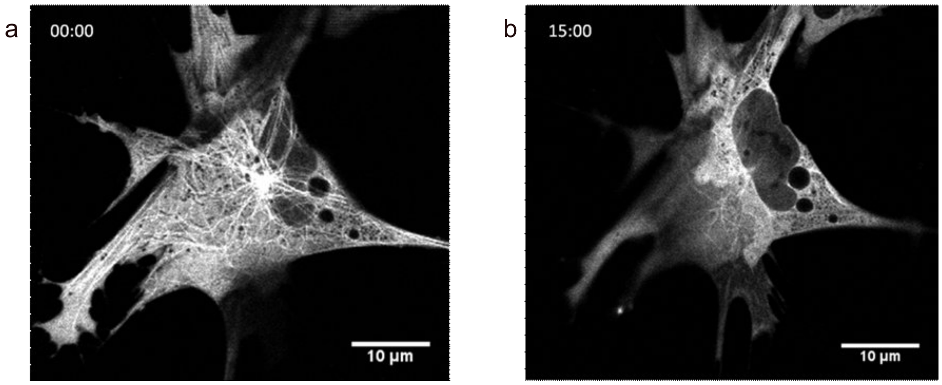


Fig. S 5. Chick fibroblast cells were treated with Nocodazole at a concentration of 16.6 µM. a) Shows intact microtubule network during initial stages of Nocodozole treatment. b) Shows that the microtubules are depolymerized after 15 minutes of the drug treatment. Tethers were pulled from these cells which were completely devoid of microtubules (see Fig. S 6 below) to check for the importance of microtubules-associated motor proteins in generating active peaks. Fibroblasts were used as complete disruption of microtubules is hard to achieve in axons without the axon undergoing severe morphological changes or atrophy

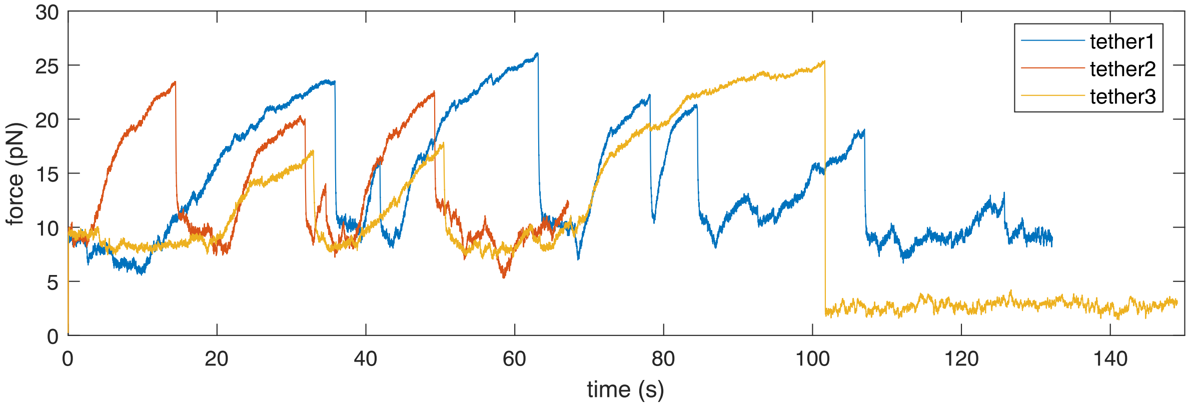


Fig. S 6. Overlayed force-time plots (data from three tethers) to show the occurrence of active peaks in presence of Nocodozole treatment on fibroblast cells.

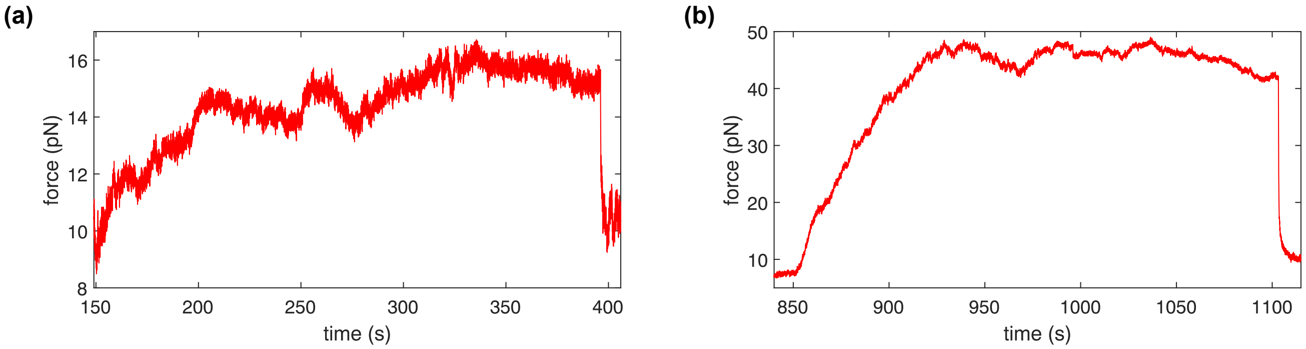


Fig. S 7. Example plots of long duration peak observed when the cells were, (a) transfected with ADF S3E (inactive mutant of ADF), and (b) transfected with SiRNA-ADF to inhibit ADF production inside cells.

#### II. MODEL

We consider a scenario where actin filaments grow inside a membrane tube held by an optical trap. As these filaments reach the end of the tube, they attach to the membrane. Within the tube, cofilin proteins bind to actin filaments, causing conformational changes, such as the shrinkage of the filaments. Due to the bound cofilins, the filaments experience forces that are subsequently transferred to the trap. As time increases, more cofilins bind to the actin filaments and exert an increasing force on the trap, eventually leading to rupture of the filaments from the trap. To explain the mechanism behind the experimentally observed peak force and rupture time distribution, we theoretically explore different detachment protocols for a *single actin filament* and compare it with the experimental data.

##### A. Helical filaments

In the main text, we consider that ADF/Cofilin binding induces the shrinkage of actin filaments which results in a pulling force at the nanotube tip (Fig. 5 a, main text). Experimental evidence of helical coiling of actin after detachment from the nanotube tip (Fig. 3 d, main text) suggest that force generation might be due to coiling rather than shrinkage. Here we show the conditions under which coiling can the represented as an effective shrinkage.

Actin filaments are semi-flexible biopolymers with an intrinsic helical structure [1] that plays a crucial role in cellular mechanics. Their mechanical properties, such as spontaneous curvature and torsion, enable them to sense and respond to external forces. In this study, the helical filament is subjected to tension as an optical trap pulls it. The mechanical properties of the filament interact with the pulling force of the optical trap, determining its optimal shape. Cofilins bind to the actin filament and induce twisting. As more cofilin bind, they generate spontaneous twists in the filament. The twisting mode couples with bending and the pulling force of the optical trap, collectively maintaining the helical structure. Here we study how cofilin-induced twists, combined with bending rigidity, reduce the end-to-end distance of the filament.

###### *1. Geometry of the helix*

The position vector to a point on helix can be expressed as

$$\mathbf{r} = R_a \hat{e}_\rho + c R_a \theta \hat{e}_z \tag{1}$$

where $R_a$ and $c$ are the radius and pitch of the helix. The unit vectors $\hat{e}_\rho$ and $\hat{e}_z$ along radial and $z$ directions similar to cylindrical coordinate system. The radial and angular unit vectors are written as

$$\hat{e}_\rho = \cos(\theta)\hat{e}_x + \sin(\theta)\hat{e}_y \quad \hat{e}_\theta = -\sin(\theta)\hat{e}_x + \cos(\theta)\hat{e}_y \tag{2}$$

The coordinate system of the helical geometry is now described using orthonormal vectors. The tangent vector $\mathbf{t}$ ($= d\mathbf{r}/ds$) aligns with the filament contour $s$, while the normal vector $\mathbf{n}$ and the binormal vector $\mathbf{b}$ ($= \mathbf{t} \times \mathbf{n}$) are given below

$$\mathbf{t} = R_a \dot{\theta} \hat{e}_\theta + R_a c \dot{\theta} \hat{e}_z, \quad \mathbf{n} = \frac{d\mathbf{t}}{ds} / \left| \frac{d\mathbf{t}}{ds} \right| = -\hat{e}_\rho, \quad \mathbf{b} = R_a \dot{\theta} \hat{e}_z - R_a c \dot{\theta} \hat{e}_\theta. \tag{3}$$

where the helix are assumed to be isotropic and homogeneous ($\ddot{\theta} = 0$). The normality condition of the vectors yields $\dot{\theta} = 1/(R_a \sqrt{1+c^2})$. In the Frenet–Serret reference frame, the curvature $K$ and the torsion $\tau$ are related by orthonormal unit vectors which are given below

$$K = \mathbf{n} \cdot \frac{d\mathbf{t}}{ds} = \frac{1}{R_a(1+c^2)}; \quad \tau = -\mathbf{n} \cdot \frac{d\mathbf{b}}{ds} = \frac{c}{R_a(1+c^2)} = cK; \tag{4}$$

This leads to

$$R_a = \frac{K}{K^2+\tau^2} = \frac{1}{K(1+c^2)} \tag{5}$$

where we use $\tau = Kc$. The force due to filament coiling is associated to the filament end-to-end distance $L_e$. Calling the (constant) filament contour length $L_c$, we define the dimensionless end-to-end distance of the filament as $\overline{\ell}_e = L_e/L_c$, which is related to the pitch by $\overline{\ell}_e = cR_a\dot{\theta} = \frac{c}{\sqrt{(1+c^2)}}$. The relation between pitch parameter $c$, $\overline{\ell}_e$, and $\tau$ which are therefore:

$$c = \frac{\overline{\ell}_e}{\sqrt{(1-\overline{\ell}_e^2)}} \quad \text{and} \quad \tau = K\,\frac{\overline{\ell}_e}{\sqrt{(1-\overline{\ell}_e^2)}} \tag{6}$$

Binding of cofilins to the actin filament induces changes in the spontaneous curvature $K_0$ and spontaneous torsion $\tau_0$ $(= K_0 c_0)$,

##### *2. Free energy of a single helical filament:*

Th elastic energy of a helical filament of spontaneous curvature $K_0$ and spontaneous torsion $\tau_0$ $(= K_0 c_0)$ can be written as

$$E_{\text{fil}} = \frac{k_B T L_c}{2}\left[\ell_b\,(K-K_0)^2 + \ell_t(\tau - K_0 c_0)^2\right], \tag{7}$$

where $L_c$ is the contour length, $k_BT$ is Boltzmann's constant times the temperature, $\ell_b$ is the bending persistence length, and $\ell_t$ is the torsional persistent length. These parameters cover a broad range for actin filament in vivo and vitro system: $\ell_b \simeq 3$ to $17\mu m$ and $\ell_t \simeq 0.5$ to $20\mu m$ [2].

Let us now consider that a trap is located $L^0_{\text{trap}}$ distance away from the base of the filament and of stiffness $K_{\text{trap}}$. In general, the trap extends the filament from its thermally equilibrated helical shape, and increase its end-to-end distance $L_e$. It is important to note that $L_e < L^0_{\text{trap}} < L_c$ where $L_c$ is the total contour length of the filament. We use dimensionless length scales: $\overline{\ell}_e = L_e/L_c$ and $\overline{\ell}_0 = L^0_{\text{trap}}/L_c$. The normalized energy of the filament per unit contour length is written as

$$\widehat{E}_{\text{fil+trap}}(K,\overline{\ell}_e) = \ell_b(K-K_0)^2 + \ell_t\left(\frac{K\,\overline{\ell}_e}{\sqrt{(1-\overline{\ell}_e^2)}} - K_0\,c_0\right)^2 + \frac{K_{\text{trap}}L}{k_BT}\,\left(\overline{\ell}_0 - \overline{\ell}_e\right)^2, \tag{8}$$

We first minimize the energy with respect to $K$ fixing $\overline{\ell}_e$, i.e., $\frac{\partial}{\partial K}\widehat{E}_{\text{fil+trap}} = 0$, and obtain

$$K^* = K_0\frac{1+\overline{\ell}_t\,c_0\frac{\overline{\ell}_e}{\sqrt{1-\overline{\ell}_e^2}}}{1+\overline{\ell}_t\frac{\overline{\ell}_e^2}{1-\overline{\ell}_e^2}}, \tag{9}$$

where $\overline{\ell}_t = \ell_t/\ell_b$. In the limit $\overline{\ell}_t \to 0$, Eq. (9) yields $K^* \to K_0$. We use Eq.9, and substitute $K = K^*$ in the expression for energy density given by Eq. 8, and minimize the energy density with respect to $\overline{\ell}_e$. This amounts to enforcing the balance of force between the trap and the helix, and yield an equation for the end-to-end length $\overline{\ell}_e$

$$\frac{f_{\text{trap}}}{K_{\text{trap}}L_c} = \left(\overline{\ell}_0 - \overline{\ell}_e\right) = \frac{K_0^2\ell_b^2\left(\overline{\ell}_e - c_0\sqrt{1-\overline{\ell}_e^2}\right)\left[\left(1-\overline{\ell}_e^2\right) + c_0\overline{\ell}_e\sqrt{(1-\overline{\ell}_e^2)}\overline{\ell}_t\right]}{\overline{K}_{\text{trap}}\,(1-\ell_e^2)\left(1-\ell_e^2+\ell_e^2\overline{\ell}_t\right)^2} \tag{10}$$

where $\overline{K}_{\text{trap}} = K_{\text{trap}}L_c/(k_BT/\ell_b)$ is a dimensionless quantity. By solving this equation numerically, we obtain the equilibrium end-to-end distance $\overline{\ell}_e^*$ of the filament and the force felt by the optical trap for given values of the geometric parameters $\overline{\ell}_0, \overline{\ell}_e$.

##### 3. *Force variation upon cofilin binding.*

The filaments grows freely before binding to the nanotube tip, so that it adopts its preferred curvature $K_0^{\text{intr}}$ and pitch $c_0^{\text{intr}}$, and reaches a contour length when it attaches to the tip that results in no force $f_{\text{trap}} = 0$. We assume this configuration is free of cofilin.
Cofilin binding follows a first order kinetics:

$$\rho_{\text{cofil}}(t) = \frac{\rho_{\max}}{1 + \frac{k_{\text{off}}}{k_{\text{on}}}} \left(1 - e^{(k_{\text{off}} + k_{\text{on}})t}\right) = \frac{\rho_{\max}}{1 + \frac{k_{\text{off}}}{k_{\text{on}}}} \hat{\epsilon}(t) \tag{11}$$

where $0 \leq \hat{\epsilon} \leq 1$. We assume that cofilin binding induces change in spontaneous curvature and twist, that can be written at first order as

$$K_0(\hat{\epsilon}) = K_0^{\text{intr}} \left(1 + a\,\hat{\epsilon}\right), \quad \tau_0(\hat{\epsilon}) = \tau_0^{\text{intr}} \left(1 - b\,\hat{\epsilon}\right), \quad c_0(\hat{\epsilon}) = \frac{\tau_0^{\text{intr}}}{K_0^{\text{intr}}} \left(\frac{1 - b\,\hat{\epsilon}}{1 + a\,\hat{\epsilon}}\right), \tag{12}$$

where $a$ and $b$ are parameters of the first order expansion.

We present numerical results for the evolution of the force with $\hat{\epsilon}$ in Fig. S8. It shows a linear relationship as the shrinkage model postulates, for a wide range of parameters. We present the numerical results for the variation of the force with time (shown in Fig. S9), assuming and first order binding kinetics for cofilin, which presents the linear growth followed by an exponential saturation and postulated by the shrinkage model. In summary, the shrinkage model with effective parameters is apt at mimicking the case where the pulling force results from a torsional stress induced by cofilin of actin filaments.

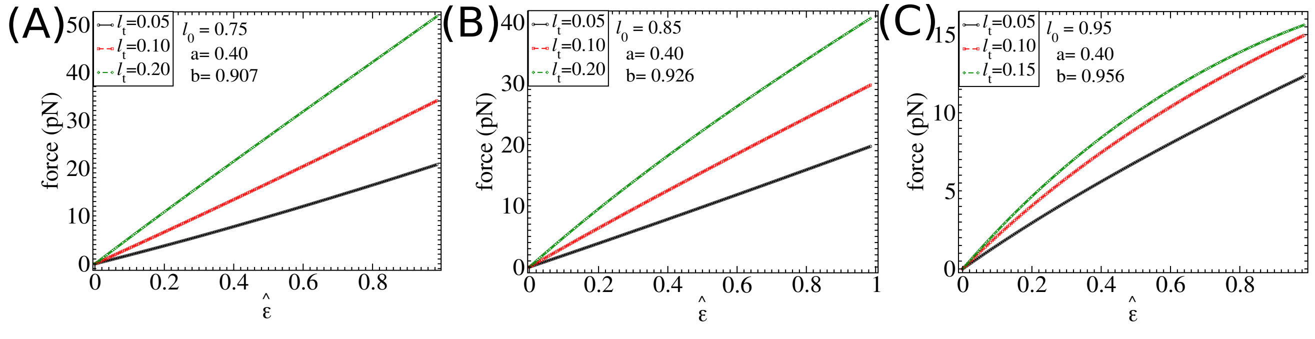


Fig. S 8. Study of force variation with the amount of bound cofilins $\hat{\epsilon}$ where we vary the ratio of the torsional and bending persistence lengths denoted as $\bar{\ell}_t$. We fix $\ell_b = 10\mu m$, and vary torsional persistence length and the value of cofilin binding rate is $k_{\text{b}} = 0.25$ in $s^{-1}$. We fix the rate of change of spontaneous curvature $a = 0.40$ given in Eq. (12) and we also consider that the end-to-end distance is reduced by 90% of the initial end-to-end distance before bound cofilins. In figure (A), (B), and (C), we show for initial ratio are $\bar{\ell}_0 = 0.75$, $\bar{\ell}_0 = 0.85$, and $\bar{\ell}_0 = 0.95$, respectively.

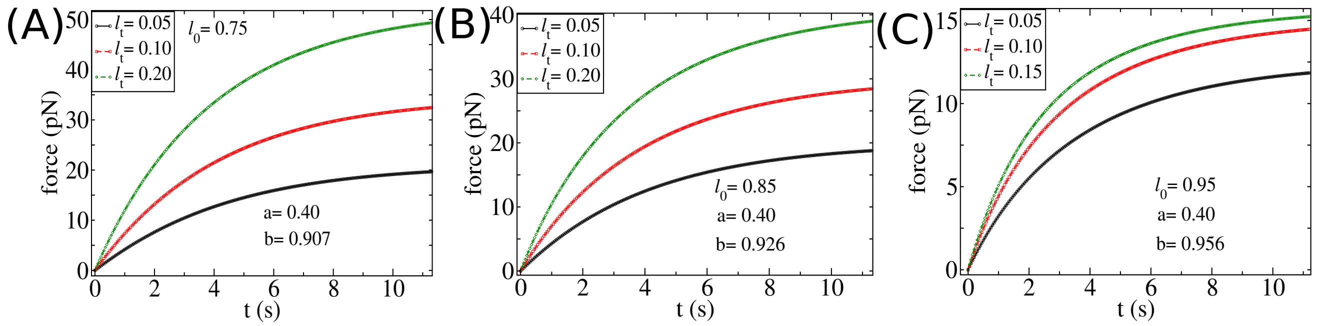


Fig. S 9. Study of force saturation with time $t$ where $\hat{\epsilon}$ is related with time is given by $\hat{\epsilon} = 1 - e^{-k_{\rm b}\, t}$. We vary the ratio of the torsional persistence length and bending persistence length denoted as $\bar{\ell}_t$ where we fix $\ell_b = 10\mu m$. The value of cofilin binding rate is $k_{\rm b} = 0.25$ in $s^{-1}$. We fix the rate of change of spontaneous curvature $a = 0.4$ given in Eq. (12) and considering end-to-end distance is reduced by 90% of the initial length.

#### B. Rupture force distribution for a single filament

##### 1. *Mechanosensitive detachment*

In the main text, we describe the time evolution of the cofilin-induced force using a single-filament model (Equation (2), main text). In the limit $N \to \infty$ (large filament length), the probability that the filament has not detached (survival probability) after a time $t$ satisfies

$$\frac{d}{dt} P_{\rm surv}(t) = -k_{\rm r} P_{\rm surv}(t) \tag{13}$$

where $k_{\rm r}$ is the rupture rate. We express $P_{\rm surv}$ in terms of the force as:

$$\frac{\partial P_{\rm surv}}{\partial f_{\rm a}} = -k_{\rm r} P_{\rm surv}/\dot{f}_{\rm a} \tag{14}$$

Using Eq.(2) in the main text, we substitute $\dot{f}_{\rm a}$ in the above expression and obtain

$$\frac{\partial P_{\rm surv}}{\partial f_{\rm a}} = -P_{\rm surv} \frac{k_{\rm r}(f_{\rm a})}{k_{\rm b}(f_{\max} - f_{\rm a})}. \tag{15}$$

We use the initial condition $P_{\rm surv}[f = 0] = 1$ and obtain

$$P_{\rm surv}[f] = \exp\left[ -\int_0^f df' \frac{k_{\rm r}[f']}{k_{\rm b}(f_{\max} - f')} \right] \tag{16}$$

##### 2. *Mechanosensitive detachment rate: Bell model*

We study the rupture event by assuming that the bond between actin and linker protein is a slip bond, following the typical Bell model for the unbinding rate: $k_{\rm r} = k_{\rm r0} e^{f_{\rm a}/f^*}$. This formulation allows us to express the survival probability as

$$P_{\rm surv}(f) = \exp\left\{ \left[ -\frac{k_{\rm r0}}{k_{\rm b}} f^* e^{f_{\max}/f^*} \left( \Gamma[0, (f_{\max} - f)/f^*] - \Gamma[0, f_{\max}/f^*] \right) \right] \right\} \tag{17}$$

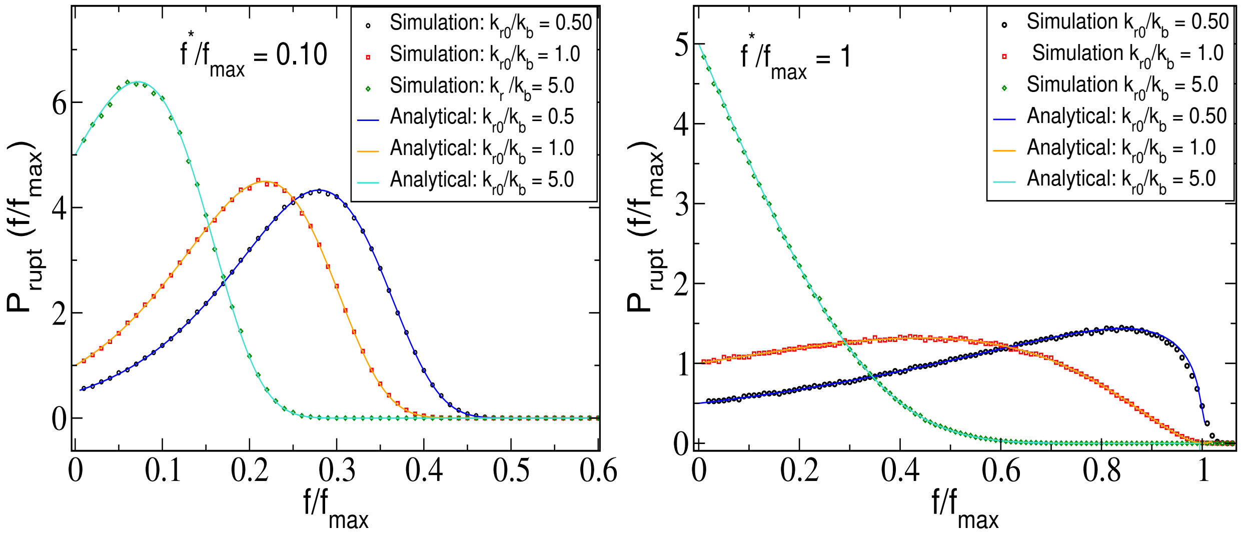


Fig. S 10. Single filament rupture force distribution. We compare the analytical results for rupture-force distribution and the simulation results for peak-force distribution for different values of parameters.

where $\Gamma[0, z]$ is the incomplete Euler Gamma function. From survival probability, one may obtain the probability density of the rupture force as $P_{\rm rupt}(f) = -\partial_f P_{\rm surv}(f)$ which can be expressed as

$$P_{\rm rupt}(f) = P_{\rm surv}(f) k_{\rm r}(f)/\dot{f} = \frac{k_{\rm r0}}{k_{\rm b}} P_{\rm surv}(f) \frac{e^{f/f^*}}{f_{\rm max} - f}. \tag{18}$$

where we have used $f(t) = f_{\rm max}\left(1 - e^{k_b t}\right)$ obtained in the main text (Eq.2).

We compare the results from the numerical simulation with Eq. S18 for different values of the parameters $f^*/f_{\rm max}$ and $k_{\rm r0}/k_{\rm b}$ shown in Fig. S10. In the strong force-sensitive limit ($f^* \ll f_{\rm max}$), the distribution exhibits a peak followed by a decay over a force scale comparable to the peak. The peak shifts to lower forces as $k_{\rm r0}/k_{\rm b}$ increases (Fig. S10-left). With increasing $f^*$, for example $f^* = f_{\rm max}$, the distribution ressembles the force-insensitive detachment probability for larger values of $k_{\rm r0}/k_{\rm b}$, and is rather flat for small values of value of $k_{\rm r0}/k_{\rm b}$ (see Fig. S10-right).

As we described before, the peak force distribution in the experiment exhibits two features: (i) a peak at small forces and (ii) exponential decay at large forces. To get the exponential decay, a constant detachment rate is required at large forces, while a peak requires Bell-like mechanosensitivity at small forces. To obtain an exponential decay, we first assume a constant detachment rate ($k_{\rm r} = k_{\rm r0}$), and vary its values to match the tail of the distribution. Though the model matches with the tail, it fails to match the rise and the location of the peak at small forces (black curve in Fig. S11-Left). Next, we assume the Bell-like detachment rate (i.e., $k_{\rm r} = k_{\rm r0}\, e^{f/f^*}$) and vary both $k_{\rm r0}$ and $f^*$. The model captures the behavior at small forces but fails to capture the slow decay at large forces (green curve in Fig. S11-Left.)

##### *3. Mechanosensitive detachment and large force saturation*

Since a simple Bell-like kinetics does not simultaneously capture the small-force and large-force behaviors of the experimental probability distribution, a modified model may be analyzed combining Bell-like detachment at small forces and a constant detachment rate at large forces, which may explain the exponential decay. The new form of the detachment rate is chosen to be:

$$k_{\rm r}(f) = k_{\rm r0} \exp\left[\frac{f f_{\rm th}}{f^*(f_{\rm th} + f)}\right] \tag{19}$$

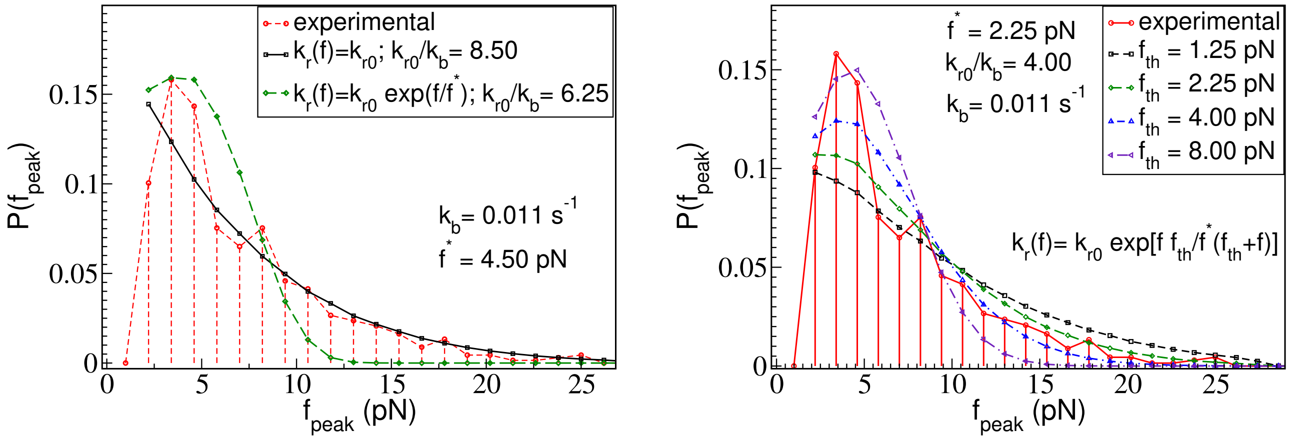


Fig. S 11. **Left**: Comparison with experiments for the peak-force distribution of Bell-like and constant detachment rates for a single filament. The figure illustrates how different models for a single filament compare with experimental observations. The black curve, which represents a constant detachment rate (same for all forces), agrees well with the tail of the distribution but does not capture the initial peak. On the other hand, the Bell-like mechanosensitive detachment rate matches the initial rise but decays fast, and fails to produce long tail of the experimentally observed peak-force distributions. **Right**: Peak-force distribution with a Bell-like force-dependent detachment rate for small forces and saturates to a constant value for large forces. The parameters $f^*$ and $k_\mathrm{r}$ are set to values that allow for the best fit, and the parameter $f_\mathrm{th}$ is varied so as to reproduce the small force or large force behaviours. For very small values of $f_\mathrm{th}$, the detachment rate is effectively constant, while for large values of $f_\mathrm{th}$, it becomes mechanosensitive.

with a force threshold $f_\mathrm{th}$ beyond which the detachment rate saturates. To fit the experimental observations, we vary $f_\mathrm{th}$ and $f^*$ in our simulation. A small value of the threshold force reproduces the results of the constant detachment rate model; an exponential tail but no peak at small forces. As $f_\mathrm{th}$ increases, a peak emerges at small forces but the slow exponential decay at large force is no longer observed. Fig. S11-Right shows that the experimental data are not well matched by this more complex single-filament model.

#### C. Model for Multiple filaments

In the experimental force-time plot (shown in Fig. S12(top panel)), we observe frequent sharp drops in force that do not lead to complete rupture. This behavior suggests that multiple filaments may be attached at any given time. Assuming each filament attaches with zero tension, we propose a mechanism of rupture events with multiple filaments which is described in the main text. In the next section, we study the rupture events with multiple filament via numerical simulations. We use the Gillespie algorithm [6] to model the binding and unbinding of cofilins as well as attachment and detachment of the filaments. Considering the agent-based processes, we incorporate the stochasticity of the reactions in the individual level and all the reactions occur in real time based on their chosen rates which is described in the main text.

We compute the peak force and rupture time distribution and compare them with the experimental observation reported in Fig.(1) in the main text. As this initial number of bound filament is unknown, the numerical simulation start with probabilities $p_i$ for having a number $i$ of cofilin-free filament attached. Filaments detach and attach at rates $k_\mathrm{r}$ and $k_\mathrm{a}$, and the simulation proceeds until detachment of all filaments, and record the time until full detachment and the peak force that was reached until then. This constitute an element of our statistical ensemble, from which we compute the peak force and rupture time distribution. To reduce the number of parameters, we assume that the stiffness of the linker, $k$, is much greater than $K_\mathrm{trap}$. In addition to that, there are several kinetic processes, such as the unbinding ($k_\mathrm{off}$) and binding ($k_\mathrm{on}$) of cofilins, and the detachment ($k_\mathrm{r0}$) and attachment ($k_\mathrm{a}$) of filaments. Note that $1/k_\mathrm{b}$ sets the timescale over which the cofilin-induced force in a single filament reaches saturation (given in Eq.(2) in the main text). From the experimental force-time plots, we often see that detachment occurs before the force reaches a saturated value. This suggests that $k_{\mathrm{r0}}$ may be of the order of $k_{\mathrm{b}}$. While we include all these processes, we simplify the model by setting a single rate, $k_{\mathrm{r0}} = k_{\mathrm{a}} = k_{\mathrm{b}}$, where $k_{\mathrm{b}} = k_{\mathrm{on}} + k_{\mathrm{off}} = 2k_{\mathrm{on}}$, assuming $k_{\mathrm{on}} = k_{\mathrm{off}}$. Consequently, we have two explicit parameters: $k_{\mathrm{on}}$ and the shrinkage of a filament $\delta\ell$ due to the binding of a single cofilin. We adjust these two parameters to match the peak force and rupture-time distributions.

#### D. Comparison between the theory and experiments

##### 1. *Force-time plots*

The measurement of forces with time due to the cofilins binding mainly exhibits three distinct features such as (i) partial ruptures, (ii) changes in slope, and (iii) saturation of force as well as the combination of these features. We compare the force-time plots from the simulation of the theoretical model with the experimental data to illustrate these three features in Figs. S12, S13, and S14 which show that the slope of the force change over time can be significantly large or small even in the absence of partial ruptures, as observed in both experiments and simulations. In addition to that, fig. S14 demonstrates how the force saturates over time before the final rupture, which is also observed in both theory and experiments. We can observe a good apparent match between experiments and simulations. Finally, in Fig. S15, we show the force-time plot for many rupture events where exist these three features along with their combinations.

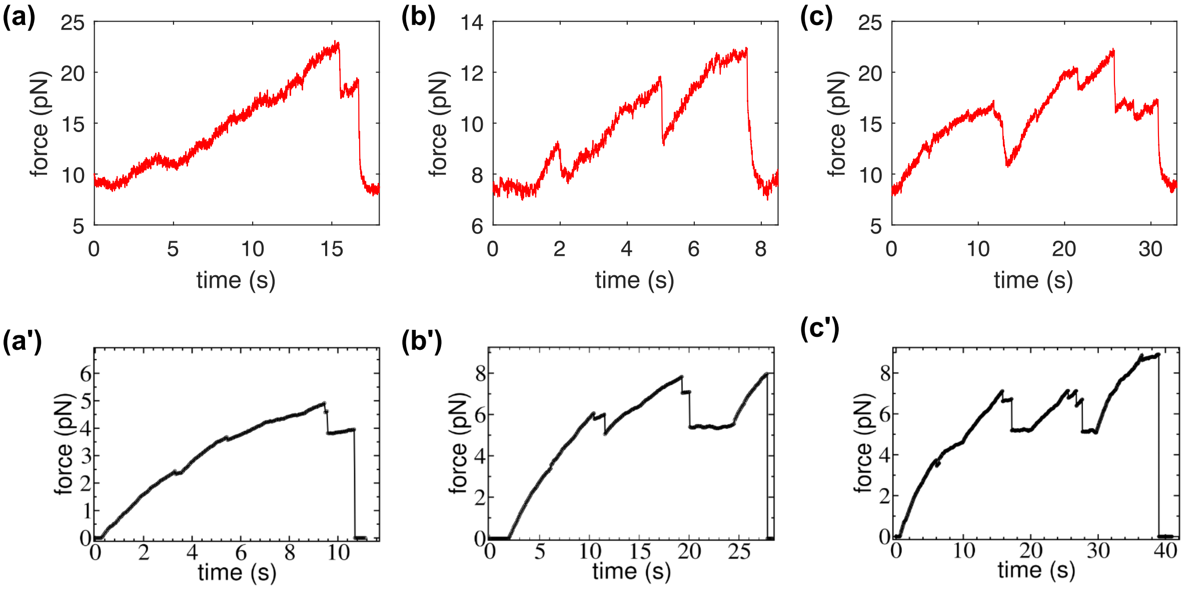


Fig. S 12. The figures show how the force decreases through *partial rupture*, followed by a rupture event, as shown in three plots. We compare the simulation (bottom panel) with the experimental (top panel) partial rupture events.

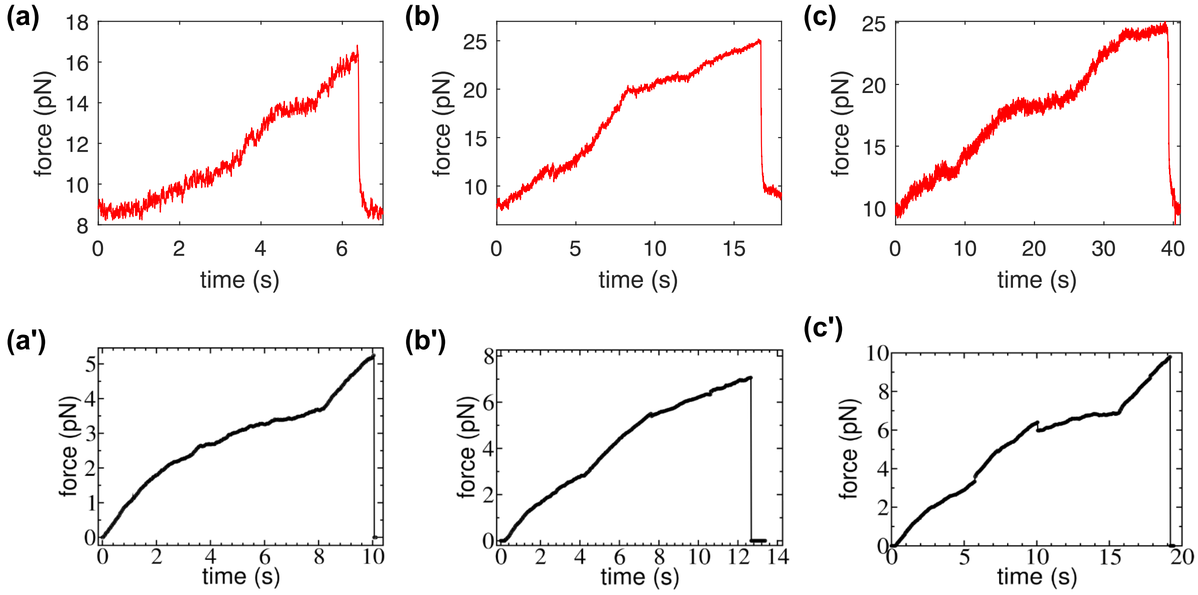


Fig. S 13. The figures show how the slope of the force-time plot changes over time. We compare the simulation (bottom panel) with the experimental data (top panel).

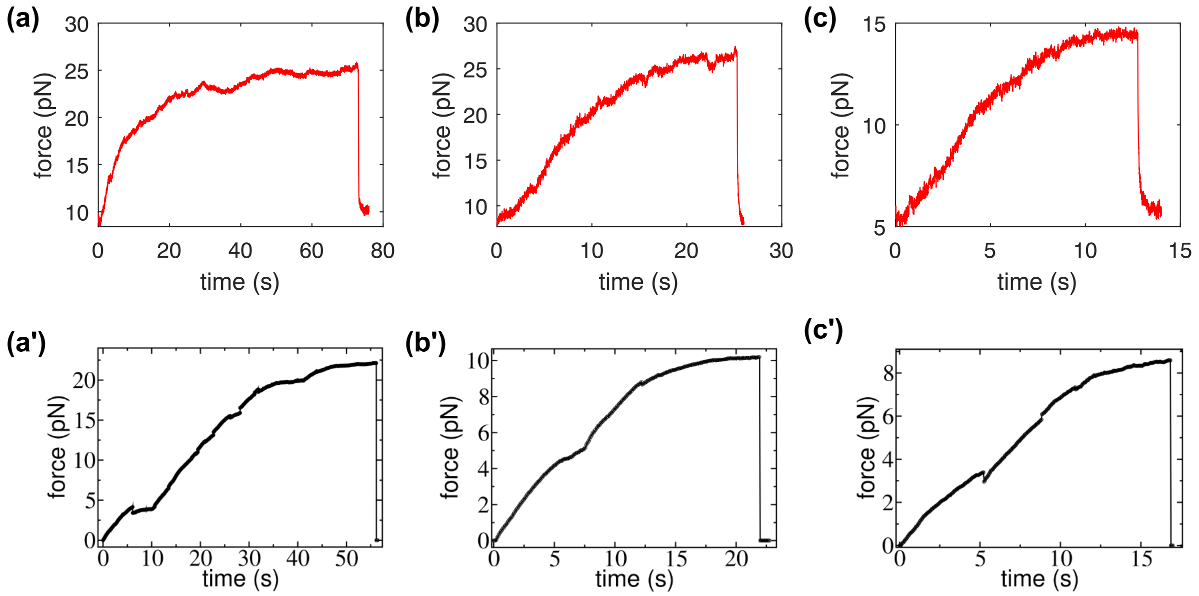


Fig. S 14. The figures show the initial development of the force and eventual saturation shown by three plots. Here we also compare the simulation with the experiment wherein the top and bottom panels correspond to experiment and simulation, respectively.

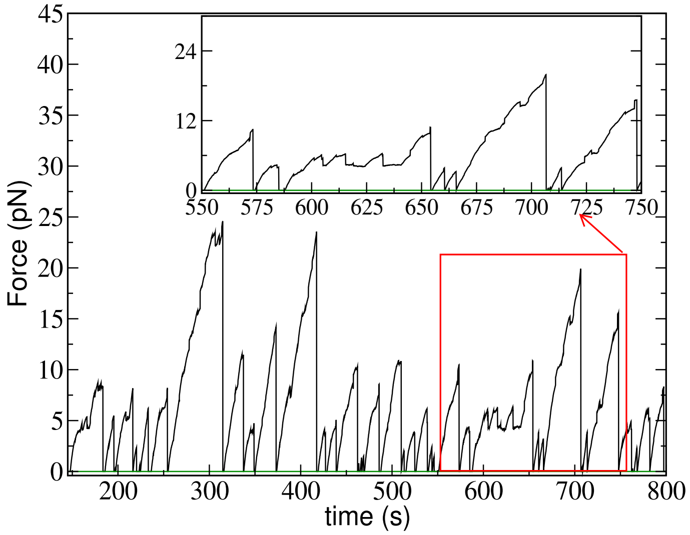


Fig. S 15. Force-time plot is shown from numerical simulation based on theoretical force-sharing model. The stiffness of the trap and length of the tether are 90pN/$\mu$m and 10$\mu$m, respectively.

#### *2. Peak force and rupture time distributions*

The experimental data for peak force and rupture time distributions show a peak at small forces and short times, respectively, followed by exponential tails. Simulations with a small number of filaments $n_{\text{fil}} = 2$ do not match the experimental observation well, in particular for the detachments at large forces. Further studies with $n_{\text{fil}} > 4$ showed that the peak of the time distribution moved to a longer times compared to experimental data. Since reattachment of filament is also considered, we found that the minimum number of filaments required to fit the experimental results is $n_{\text{fil}} = 3$. The next important parameter is the number of those filaments that are attached when the force starts rising ($t = 0$ in the simulations). We define $p_i$ as the probability of having $i$ filaments attached initially. Fig. 16 presents the results for the cases where one ($p_1 = 1$), two ($p_2 = 1$), or three ($p_3 = 1$) filaments attached. A good match to the data requires more than one filament attached initially. We argue that at the onset of each new force evolution, multiple filaments can attach with probabilities $p_1 = 3/7$, $p_2 = 3/7$, and $p_3 = 1/7$, assuming all filaments ($n_{\text{fil}} = 3$) are identical.

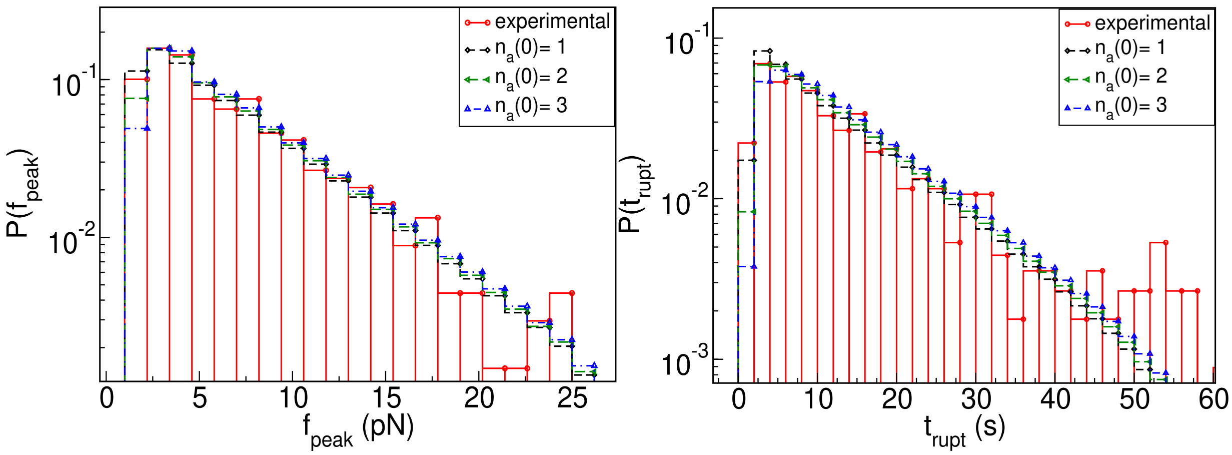


Fig. S 16. The peak force and rupture time distribution starting with one ($n_{\rm a}(0) = 1$), two ($n_{\rm a}(0) = 2$), or three ($n_{\rm a}(0) = 3$) filaments with probability 1. Note that total number of filaments is 3, the number of filaments which are initially attaches are varied.

---

[7] N. G. van Kampen, *Stochastic Processes in Physics and Chemistry, Elsevier Science Publishers, Amsterdam, (1992)*
